# Activation of basal forebrain astrocytes induces wakefulness without compensatory changes in sleep drive

**DOI:** 10.1101/2023.01.09.523360

**Authors:** Ashley M. Ingiosi, Christopher R. Hayworth, Marcos G. Frank

**Author notes:** Corresponding author (MGF).

## Abstract

Mammalian sleep is regulated by a homeostatic process that increases sleep drive and intensity as a function of prior wake time. Sleep homeostasis has traditionally been thought to be a product of neurons, but recent findings demonstrate that this process is also modulated by glial astrocytes. The precise role of astrocytes in the accumulation and discharge of sleep drive is unknown. We investigated this question by selectively activating basal forebrain (BF) astrocytes using designer receptors exclusively activated by designer drugs (DREADDs). Activation of the G_q_-protein-coupled pathway in BF astrocytes produced long and continuous periods of wakefulness that paradoxically did not cause the expected homeostatic response to sleep loss (e.g., increases in sleep time or intensity). Further investigations showed that this was not due to indirect effects of the ligand that activated DREADDs. These findings suggest that the need for sleep is not driven by wakefulness per se, but specific neuronal-glial circuits that are differentially activated in wakefulness.

## Introduction

An evolutionarily conserved feature of sleep is that it is regulated by two major processes. A biological clock provides timing (circadian) signals that organize sleep and wakefulness across the 24-hour day. A homeostatic mechanism increases sleep drive and (in some species) intensity as a function of prior wakefulness. Great progress has been made characterizing the circadian regulation of sleep and wakefulness. For example, the anatomical location of a principal biological clock and the key molecular mechanisms of timekeeping are known [1]. In contrast, far less is known about the cellular basis of sleep homeostasis. No master mammalian homeostat has been discovered and comparatively little is known about the molecular cascades in brain cells necessary for sleep homeostasis. Nevertheless, until recently, the traditional view has been that sleep homeostasis is a product of neurons.

We showed that sleep homeostasis is also dependent on glial astrocytes [2,3]. Astrocytes perform many critical functions in the brain that make them well-positioned to mediate sleep homeostasis [4]. For example, astrocytes express receptors for wake-promoting neuromodulators like noradrenaline [5,6], and astrocytes release sleep-promoting substances like ATP/adenosine which alters neuronal activity in canonical sleep-wake nuclei [7-9]. In addition, conditional inhibition of gliotransmission in astrocytes inhibits sleep drive, as measured by sleep-wake behavior and electroencephalograph (EEG) activity [7]. Under these conditions, transgenic mice show less compensatory responses to sleep loss, which suggests that they can stay awake with less accumulating sleep drive. Conditional deletion of astrocytic membrane-bound receptors to circulating neuromodulators [2] or intracellular calcium regulating proteins produces similar effects [3]. This suggests that astrocytes respond to extracellular signals (ligands) originating from neurons (e.g., classic neurotransmitters or neuromodulators).

To explore this further, we investigated a key pathway that links membrane-bound receptors to secondary intracellular cascades in astrocytes. More specifically, mammalian astrocytes express *in vivo* several membrane-bound receptors that couple to G-proteins. Therefore, experimentally manipulating these pathways engages native mechanisms present *in vivo*. This can be accomplished by using selective astroglial expression of designer receptors exclusively activated by designer drugs (DREADDs) coupled to the G_q_-protein pathway. In astrocytes, G_q_ activates the phospholipase C pathway which increases intracellular calcium through inositol 1,4,5-trisphosphate-mediated release from internal stores [10-13]. Muscarinic acetylcholine receptors [11], group I metabotropic glutamate receptors [14], and histamine H_1_ receptors [15] are expressed by astrocytes and trigger G_q_-protein coupled cascades. Research on the impact of astroglial G_q_-DREADD activation on sleep expression is limited, but recent studies show the effects are region-specific. G_q_-DREADD activation of cortical astrocytes increases non-rapid eye movement sleep (NREMS) duration [10] whereas astroglial activation in the lateral hypothalamus increases wakefulness [16]. However, the impact of astroglial DREADD activation on sleep homeostasis has yet to be explored.

We examined the effects of chemogenetically activating the G_q_ pathway in basal forebrain (BF) astrocytes. This is because the BF is comprised of several classes of neurons that influence sleep and wake time as well as sleep homeostasis [17,18]. Furthermore, the role of BF astrocytes in sleep expression and regulation has not been investigated. We find that activating G_q_-DREADDs in BF astrocytes leads to hours of sustained wakefulness without the normal compensatory changes in sleep time or intensity.

## Results

### CNO activation of astroglial Gq DREADDs in BF promotes sustained wakefulness

We first determined the impact of activating BF astrocytes on sleep-wake architecture and EEG activity. To do this, we expressed G_q_ DREADDs selectively in astrocytes by crossing Aldh1l1-Cre^+/-^ mice with hM3Dq^fl/-^ mice to produce Aldh1l1-Cre^+/-^; hM3Dq^fl/-^ (G_q_^+/-^) mutant offspring and Aldh1l1-Cre^-/-^; hM3Dq^fl/-^ control (Ctrl) littermates (S1 Fig). We then activated astroglial G_q_ DREADDs by delivering the DREADD ligand clozapine N-oxide dihydrochloride (CNO) directly to the BF. We adopted procedures from Adamsky *et al* [19] to verify CNO increased activity in BF astrocytes. As shown in S1 Fig, CNO in G_q_^+/-^ mice significantly increased cFos expression in BF astrocytes compared to CNO in Ctrl mice.

We found that activating BF astroglial G_q_ DREADDs in G_q_^+/-^ mice induced long periods of continuous wakefulness (≥ 6 h) and an overall increase in wake time for the entire light phase (Figs 1 and 2A, S1 Table) when CNO was injected during Zeitgeber time (ZT) 0. This was accompanied by increased latencies to sleep (S2A Fig, S2 Table) as well as reduced sleep time, bout frequency, and bout duration for NREMS and rapid eye movement sleep (REMS) for the entire 12-h light period (Fig 2, S1 Table). During the dark period, G_q_^+/-^ sleep was more fragmented after CNO compared to vehicle as reflected by shorter, more frequent bouts of wakefulness and NREMS. Dark period sleep time after CNO did not differ from vehicle values except for a transient increase in NREM sleep time in the first 2 h of the dark phase (h13 – 14; Fig 2).

**Fig 1:**
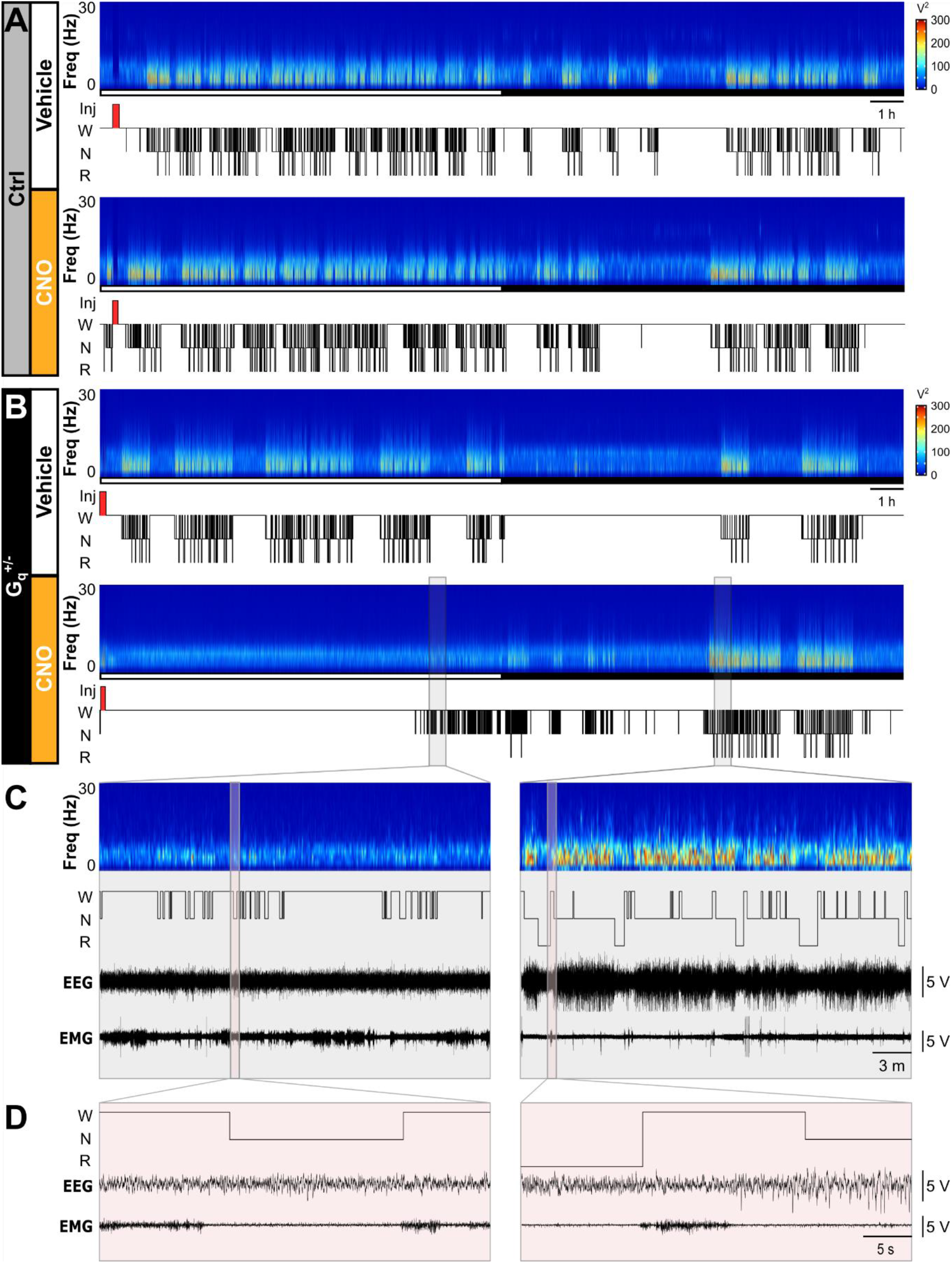
CNO activation of G_q_-DREADDs in basal forebrain astrocytes promotes wakefulness. Representative spectrograms, hypnograms, and EEG &EMG traces show responses to vehicle or CNO injections to the BF from a (**A**) Ctrl mouse and a (**B**) G_q_^+/-^ mouse. Open and closed bars below spectrograms represent the light and dark periods, respectively. (**C**) Gray boxes show 30-min subsets of G_q_^+/-^ post-CNO injection data from B for low power, ‘fragmented’ sleep (left) and ‘recovered’ sleep (right). (**D**) Red boxes show 40-s subsets of G_q_^+/-^ post-CNO injection data extracted from the gray boxes in C. Voltages are ×10^5^. Injection timepoints are highlight by red rectangles in the hypnogram. Freq, frequency; Inj, injection; W, wakefulness; N, NREMS; R, REMS.

**Fig 2:**
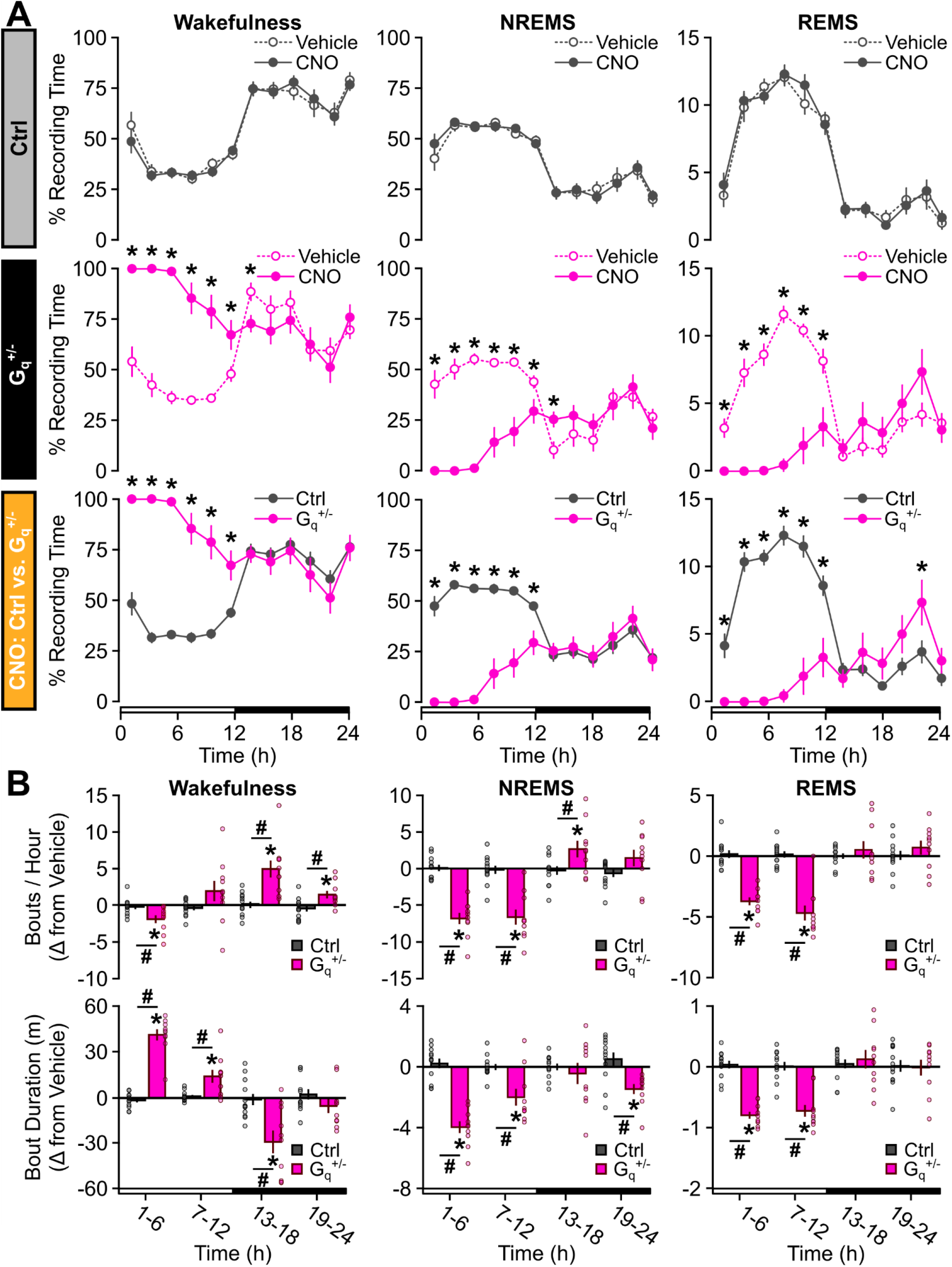
G_q_^+/-^ mice show prolonged, consolidated wakefulness and reduced sleep after CNO. (**A**) Time spent in wakefulness (left), NREMS (middle), and REMS (right) after vehicle or CNO delivery to BF during ZT0 shown as a percentage of total recording time in 2-h bins for Ctrl (top) and G_q_^+/-^ (middle) littermates. Bottom row compares Ctrl and G_q_^+/-^ responses to CNO (repeated measures ANOVA; * p < 0.05). (**B**) Change in bout frequency (top) and bout duration (bottom) shown as CNO - vehicle differences in 6-h bins for wakefulness (left), NREMS (middle), and REMS (right). Open and closed bars on the x-axis represent the light and dark periods, respectively. *, different from 0 (one-sample t test). ^#^, different from Ctrl (repeated measures ANOVA). Values are means ± SE from n = 12 Ctrl and n= 10 G_q_^+/-^ mice. Dots in B are data from individual mice. p < 0.05.

We then verified that CNO had no effects on sleep architecture in Ctrl mice. This is a necessary control when using CNO as it has been shown to metabolize to compounds (e.g., clozapine) that can indirectly impact sleep [20-24]. The same dose delivered to Ctrl littermates had no impact on sleep architecture compared to vehicle (Figs 1 and 2, S1 Table). In addition, between-subjects comparisons of Ctrl vs. G_q_^+/-^ CNO responses recapitulated the within-subject vehicle vs. CNO comparisons in G_q_^+/-^ mice (Fig 2, S2A Fig).

### CNO-induced wakefulness is not associated with increased sleep drive

CNO-induced wakefulness in G_q_^+/-^ mice did not result in homeostatic and compensatory increases in sleep time, continuity, or intensity at sleep onset as would be expected after sleep deprivation (SD) by gentle handling [25]. Once G_q_^+/-^ mice fell asleep after the CNO injection, they exhibited reduced NREM delta (δ) power in the δ (0.5 – 4 Hz) band as well as the low δ (0.5 – 1.5 Hz) band which may be more sensitive to sleep loss [7] (Fig 3A, S3 Table). CNO did not affect NREM δ power of Ctrl mice (Fig 3A). NREM EEG spectral power in the low δ, alpha (α), and beta (β) frequency bands also decreased for at least 10 h post-CNO delivery in G_q_^+/-^ mice (Fig 3B, S4 Table). We then examined EEG changes in CNO-induced wakefulness to determine if other indices of increased sleep drive were also absent (e.g., waking EEG theta (θ) activity [26,27], leakage of slow waves into wakefulness [28]). We found that waking θ activity did not increase, nor did δ “leak” into waking or REMS spectra (S3A Fig, S4 Table).

**Fig 3:**
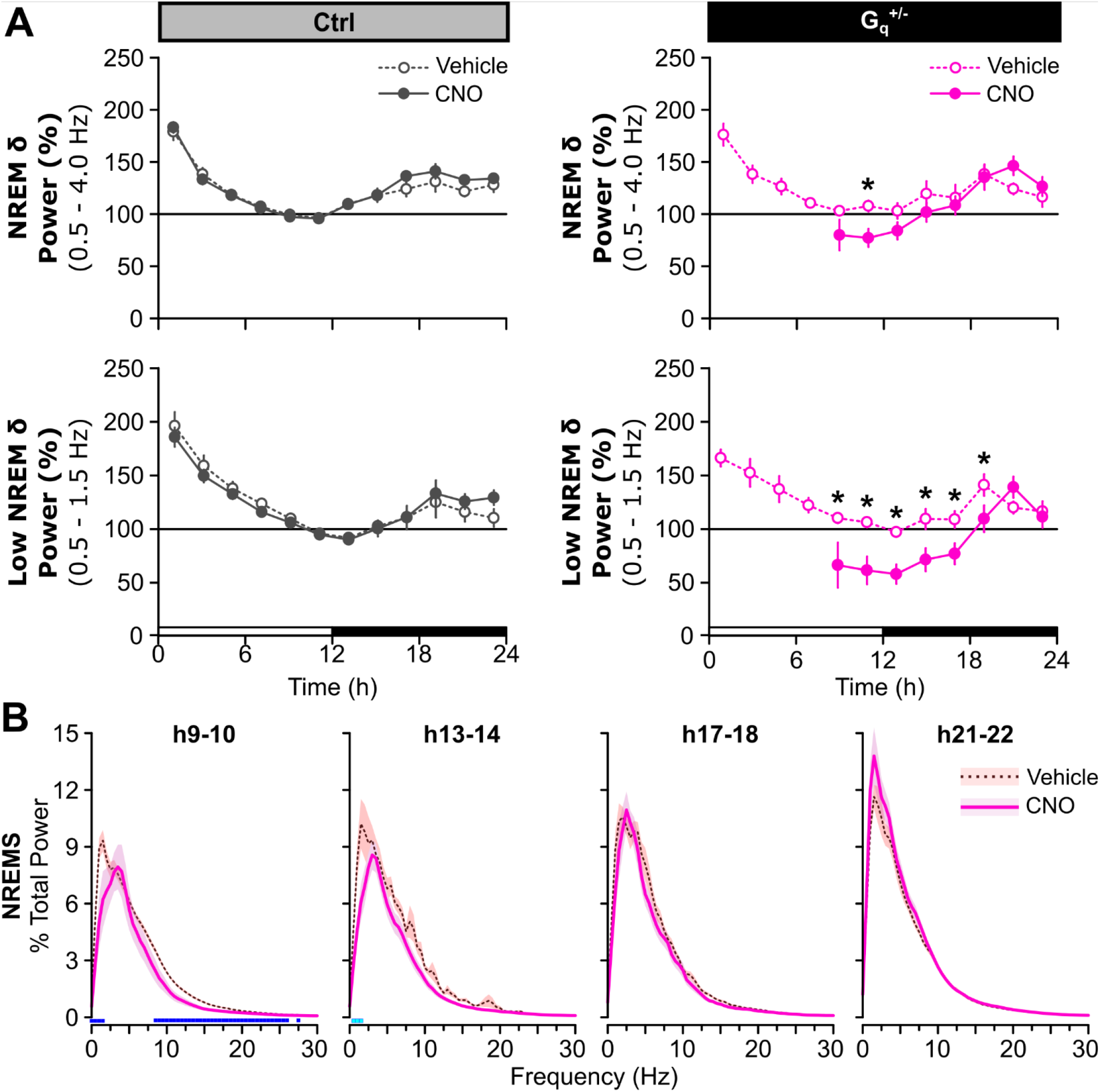
CNO does not increase sleep propensity in G_q_^+/-^ mice. (**A**) Normalized NREM delta (δ) power (0.5 – 4 Hz; top) and low NREM δ power (0.5 – 1.5 Hz; bottom) shown in 2-h bins post-injection for Ctrl (left) and G_q_^+/-^ (right) mice (Kruskal-Wallis). Open and closed bars on the x-axis represent the light &dark periods, respectively. Values are means ± SE from n = 12 Ctrl and n = 10 G_q_^+/-^ mice. * p < 0.05. (**B**) Normalized NREM EEG spectral power for G_q_^+/-^ mice shown in 2-h bins starting from the time at which (h9 - 10) ≥ 5 mice spent ≥ 5 min in NREMS within the time bin post-CNO injection. Light and dark blue squares above the x-axis denote frequency bins with significant vehicle vs. CNO differences (light blue: repeated measures ANOVA, 0.5 – 4 Hz; dark blue: repeated measures ANOVA, 0 – 30 Hz). Values are means (line) ± SE (shading). p < 0.05.

As an additional control, we measured the homeostatic response to 6 hours of SD to determine if the G_q_^+/-^ mice had a normal homeostatic response to SD (without CNO). This also provided comparison data to changes following DREADD activation in G_q_^+/-^ mice. We found that sleep homeostasis as measured by NREM EEG activity, sleep time, and sleep continuity was normal in G_q_^+/-^ mice. Six hours of SD using gentle handling produced the expected increase in NREM δ power, NREMS time, and sleep continuity (defined as fewer, but longer, NREMS bouts) during recovery compared to undisturbed baseline (BL) conditions (S4 Fig, S5 Table).

### DREADD activation: effects on body temperature and motor activity in G_q_^+/-^ mice

We assessed CNO induced changes in core body temperature and cage activity in G_q_^+/-^ mice, as large changes in these parameters might indirectly impact sleep and EEG activity [29]. CNO reduced core body temperature for ∼16 h post-injection and reduced cage activity during the dark period (S5A Fig, S6 Table). Consequently, we tested if a different DREADD ligand—JHU37160 dihydrochloride (J60)— reproduced the sleep architecture effects of CNO in G_q_^+/-^ mice (J60 does not metabolize to clozapine) [30]. J60 had similar effects on sleep architecture and EEG activity as CNO without affecting core body temperature or motor activity in G_q_^+/-^ mice (S5B Fig, S6 Table). Like CNO, J60 increased wakefulness for ∼8 h post-J60 injection in G_q_^+/-^ mice compared to vehicle (Fig 4A, S6 Fig, S1 Table) with corresponding reductions in NREM and REM sleep time, bout frequency, and bout duration (Fig 4A – B, S1 Table). J60 also increased latencies to NREMS and REMS in G_q_^+/-^ mice (S2B Fig, S2 Table). G_q_^+/-^ sleep expression in the dark period after J60 did not differ from vehicle (Fig 4).

**Fig 4:**
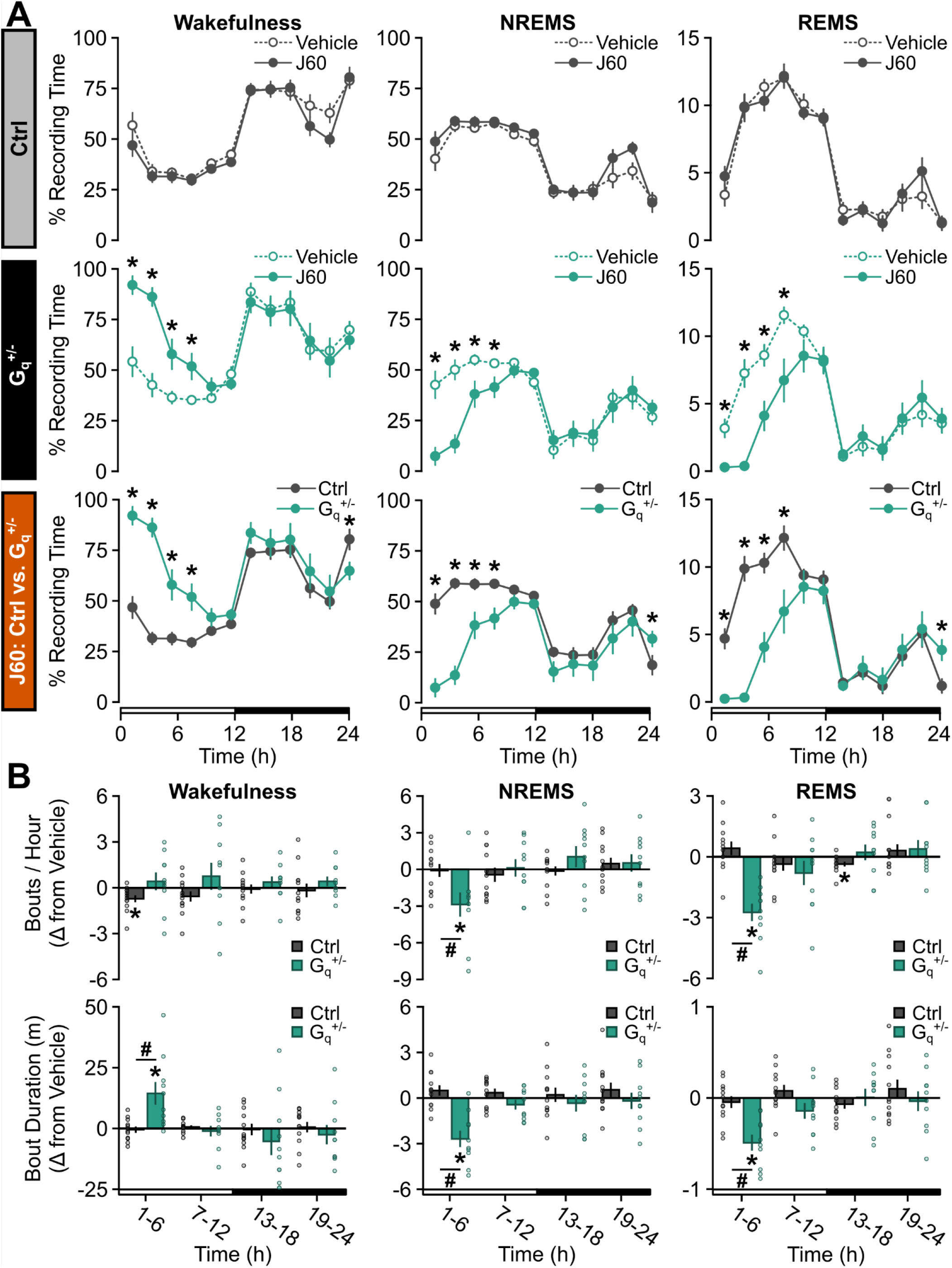
G _q_^+/-^ mice show prolonged, consolidated wakefulness and reduced sleep after J60. (**A**) Time spent in wakefulness (left), NREMS (middle), and REMS (right) after vehicle or J60 delivery to BF during ZT0 shown as a percentage of total recording time in 2-h bins for Ctrl (top) and G_q_^+/-^ (middle) littermates. Bottom row compares Ctrl and G_q_^+/-^ responses to J60 (repeated measures ANOVA; * p < 0.05). (**B**) Change in bout frequency (top) and bout duration (bottom) shown as J60 - vehicle differences in 6-h bins for wakefulness (left), NREMS (middle), and REMS (right). *, different from 0 (one-sample t test). ^#^, different from Ctrl (repeated measures ANOVA). Open and closed bars on the x-axis represent the light and dark periods, respectively. Values are means ± SE from n = 12 Ctrl and n = 10 G _q_^+/-^ mice. Dots in B are data from individual mice. p < 0.05.

Like CNO, J60-induced wakefulness did not increase sleep drive in G_q_^+/-^ mice as measured by changes in sleep time, sleep continuity, and EEG activity. As was true for CNO-induced wakefulness, J60-induced wakefulness did not lead to an increase in NREM δ power (Fig 5A – B, S3 and S7 Tables) in G_q_^+/-^ mice compared to vehicle. Similarly, J60-induced waking EEG did not exhibit elevated θ after J60, nor was there elevated δ in wake or REM spectra (S3B Fig, S7 Table).

**Fig 5:**
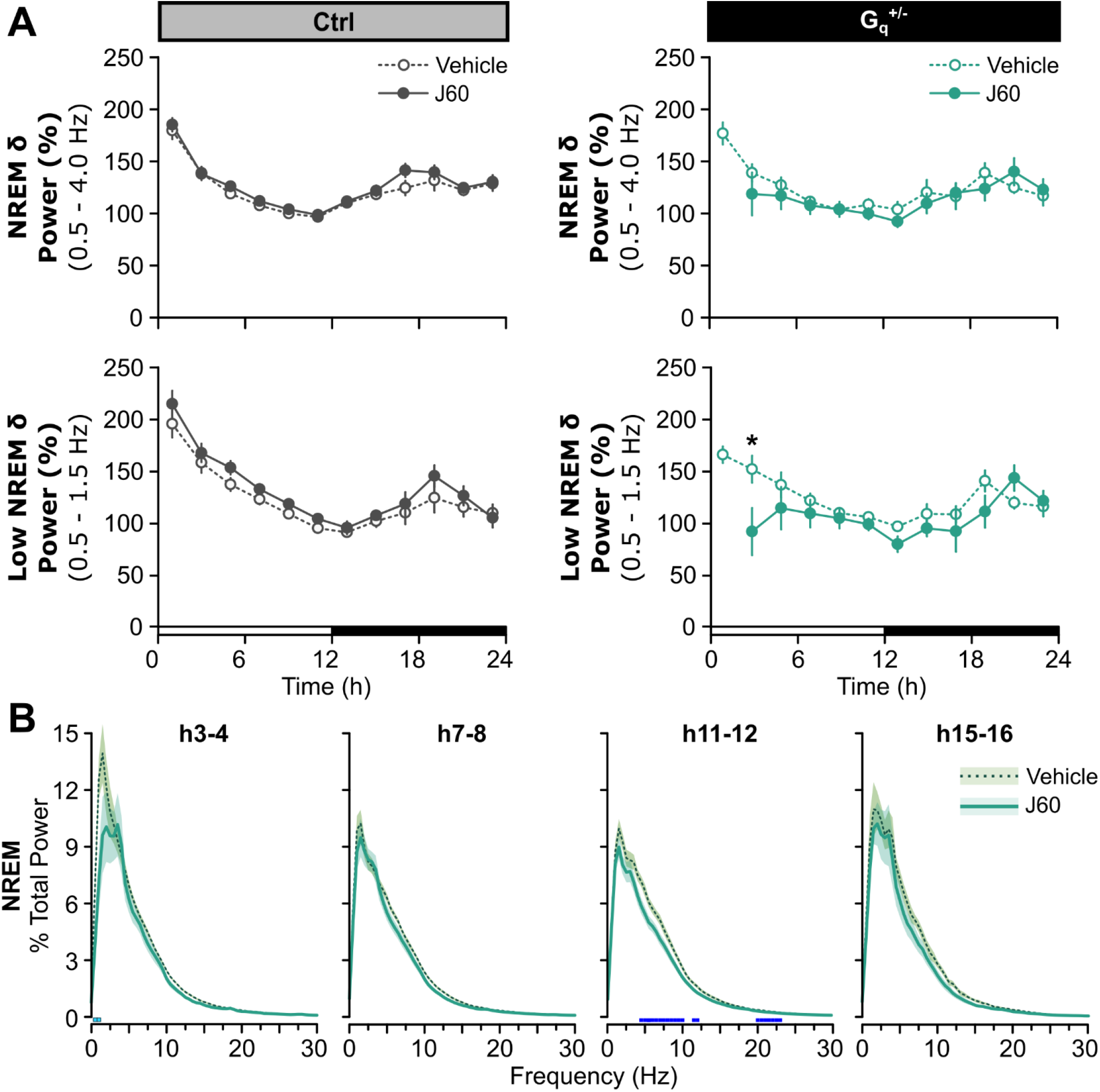
J60 does not increase sleep propensity in G_q_^+/-^ mice. (**A**) Normalized NREM delta (δ) power (0.5 – 4 Hz; top) and low NREM δ power (0.5 – 1.5 Hz; bottom) shown in 2-h bins post-injection for Ctrl (left) and G_q_^+/-^ (right) mice (Kruskal-Wallis). Open and closed bars on the x-axis represent the light &dark periods, respectively. Values are means ± SE from n = 12 Ctrl and n = 10 G_q_^+/-^ mice. * p < 0.05. (**B**) Normalized NREM EEG spectral power from G_q_^+/-^ mice shown in 2-h bins starting from the time at which (h3 - 4) ≥ 5 mice spent ≥ 5 min in NREMS within the time bin post-J60 injection. Light and dark blue squares above the x-axis denote frequency bins with significant vehicle vs. J60 differences (light blue: repeated measures ANOVA, 0.5 – 4 Hz; dark blue: repeated measures ANOVA, 0 – 30 Hz). Values are means (lines) ± SE (shading). p < 0.05.

We also found that sleep expression of Ctrl mice was mostly unaffected by J60 compared to vehicle (Fig 4 and 5, S6 Fig, S1 and 3 Tables). Between-subjects comparisons of Ctrl vs. G_q_^+/-^ J60 responses recapitulated G_q_^+/-^ vehicle vs. J60 within-subject comparisons (Fig 4, S2B Fig) as well.

### Sex differences

There were slight sex differences following different manipulations in either Ctrl or G_q_^+/-^ mice. As these did not change the overall reported effects discussed above, they are summarized in S8 – 11 Tables.

## Discussion

We investigated the role of BF astrocytes in sleep expression and regulation. We find that DREADD activation of the G_q_ pathway in BF astrocytes produces long periods of continuous waking that paradoxically do not trigger compensatory changes in sleep. Previous studies showed as little as 70 minutes of sustained wakefulness increases NREM delta power (a canonical index of sleep drive) in adult mice across diverse mouse strains [25]. In the present study, activation of BF G_q_ astroglial pathways produced ≥ 6 hours of sustained wakefulness with no compensatory changes in sleep. This finding is unlikely explained by indirect effects of the DREADD ligand CNO as these results were reproduced using a second DREADD ligand (J60) and not found in non-DREADD expressing Ctrl mice treated with CNO. We discuss our main results below in more detail.

### BF astrocytes induce waking without increasing sleep drive or intensity

DREADD G_q_ activation of BF astrocytes produced hours of waking without the expected signs of increased sleep drive. In rodents, comparable amounts of sleep loss via gentle handling, forced locomotion, or introduction of novel objects reliably results in compensatory increases in NREM EEG δ power, sleep continuity (e.g., fewer but longer NREMS bouts), and to a lesser extent, sleep time [25,31,32]. Increased sleep drive is also reported to increase waking EEG θ and δ during SD [26-28]. In contrast, despite producing waking amounts (≥ 6 hours) that lead to saturating sleep homeostatic responses in mice [25,33], CNO-induced sleep loss produced no compensatory changes in any of these metrics.

There are few examples of such complete dissociations in mammals [34]. For example, sleep deprivation does not increase NREM δ power in developing rodents until weaning. However, even in these cases, a need for sleep is clearly present, as sleep drive rapidly increases with sleep deprivation, and during recovery neonates show compensatory changes in sleep time or duration [35,36]. In addition, chronic sleep restriction fails to increase sleep time and depth even when behavioral signs of sleepiness (e.g., reduced latency to sleep) are present [37]. Although it is possible that waking experience (as opposed to being awake per se) determines adult mammalian sleep drive, this appears to be a small factor and is not always observed [34,38,39]. There are also a handful of gene mutations that influence mammalian sleep homeostasis [34,40,41], but even in conditional manipulations of these genes, sleep homeostasis is blunted [2,3,42,43], not eliminated as we show here. Previous studies reporting prolonged waking after lateral hypothalamic astrocyte activation [16] or neuronal activation (in the mammillary bodies) [44] did not examine in detail compensatory changes in subsequent sleep and EEG activity. Therefore, the latter studies are inconclusive with respect to this question.

What may then explain the unusual finding of wakefulness without sleep drive? The absence of a normal homeostatic response to sleep loss is not explained by indirect effects of CNO or an underlying defect in G_q_^+/-^ mice. For example, CNO in Ctrl mice did not alter sleep-wake architecture, and while it did decrease core temperature in G_q_^+/-^ mice, this fell within physiological ranges that occur across the sleep-wake cycle [45-47]. Moreover, this change in core temperature is an unlikely explanation for our results. This is because a second DREADD ligand (J60) that did not change core temperature or activity, reproduced the main effects of CNO on sleep time, sleep-wake architecture, and sleep homeostasis (Figs 4 and 5, S5 Fig). Sleep homeostasis was also intact in the G_q_^+/-^ mice as they responded with normal homeostatic responses to 6 h of SD (in the absence of CNO or J60). Like other mouse strains, 6 h of SD by gentle handling resulted in compensatory changes in NREM EEG δ power, sleep time, and sleep continuity during recovery (S4 Fig).

An alternative explanation is that activation of BF astrocytes changes the activity of surrounding BF neurons that play different roles in generating waking and separately, the homeostatic sleep response to waking. The BF is comprised of a heterogenous collection of neurons that may play different roles in sleep &wake, EEG activity, and the homeostatic response to sleep loss [17,48]. For example, BF cholinergic neurons (ChAT^+^) have been specifically linked to mammalian sleep homeostasis. Ablation of ChAT^+^ neurons reduces NREM compensatory responses to SD and waking EEG changes indicative of sleep drive [49,50] (but see [51]). Selective DREADD activation of BF GABAergic neurons produces long periods of wakefulness, similar to what we report, but once sleep commences, the homeostatic response is observed (e.g., increases in NREM δ power) [52]. While speculative, our findings suggest that G_q_-mediated activation of BF astrocytes leads to a complex activation of these circuits, such that waking is triggered (possibly via GABAergic activity) while cholinergic activity is inhibited (explaining the lack of a homeostatic sleep response). This explanation is also in keeping with the fact that BF astrocytes do not have long-range connections to forebrain or hindbrain canonical sleep &wake centers, which means their effects must be locally mediated by surrounding neurons that have such projections.

### Mechanisms downstream of astroglial Gq activation

Astroglial G_q_ activation triggers several downstream events that influence the activity of surrounding neurons in ways that might explain our results. These include gliotransmission, neurotransmitter uptake, and metabolic neuronal support. Of these, gliotransmission enjoys the most empirical support based on studies *in vitro, in situ*, and *in vivo*. For example, studies *in vitro, in situ*, and *in vivo* show that astroglial G_q_ activation stimulates gliotransmission of ATP [5,53-55]. ATP has diverse effects on neurons depending on their complement of adenosine or purinergic receptors. The ATP metabolite adenosine via A1 receptors can inhibit neurons or excite neurons via disinhibition (via GABAergic interneurons) [17,56-58]. Direct activation of BF purinergic receptors has also been shown to profoundly alter sleep and wake architecture [59]. Therefore, local release of astroglial ATP might be expected to have complex effects on BF neurons. In contrast, while astroglial G_q_ activation can influence neurotransmitter and ion uptake [60,61] as well as changes in astroglial-neuronal metabolism [62], there is less support for these mechanisms *in vivo*.

### Future directions

Our results raise several questions that are beyond the scope of this single study to answer. It will be important to determine changes in surrounding brain cells following astroglial BF G_q_ activation (and inhibition). Does this manipulation lead to a specific and complex activation and inhibition of BF neurons as we propose? If so, what mediates this diversity of responses? If indeed gliotransmission is the most plausible mechanism to explain how changes in astrocytes result in neuronal changes (which is yet to be determined), what are the roles of other putative gliotransmitters that have been shown to influence sleep &wake architecture and regulation? For example, G_q_-DREADD activation of astrocytes induces release of the excitatory neurotransmitter glutamate [11,63], and chemogenetic activation of BF neuronal subtypes produces different components (e.g., cholinergic-induced suppression of EEG spectral power across states) [52] of the astroglial-induced phenotype described here. Alternatively, BF astrocytes may be a heterogeneous population whose activation can result in differential downstream impacts on BF neurons. And importantly, what are the implications of producing wakefulness without cost? There are at least two implications worth discussing. The first is that this narrows the search for the need for sleep and by extension, sleep function [64]. The second is that it holds the promise of creating waking brains that need less sleep.

## Materials and methods

### Animals

B6;FVB-Tg(Aldh1l1-cre)JD1884Htz/J (Aldh1l1-Cre; 023748) and B6N;129-Tg(CAG-CHRM3*,-mCitrine)1Ute/J (hM3Dq; 026220) mice were obtained from The Jackson Laboratory (Bar Harbor, ME, USA). Heterozygous Aldh1l1-Cre^+/-^ male mice were bred with heterozygous hM3Dq^fl/-^ female mice to produce Aldh1l1-Cre^-/-^; hM3Dq^fl/-^ control (Ctrl) mice and Aldh1l1-Cre^+/-^; hM3Dq^fl/-^ (G_q_^+/-^) experimental littermates. Mice were housed in standard cages at ambient temperature 24 ± 1°C on a 12:12 h light:dark cycle with food and water *ad libitum*. All experimental procedures were approved by the Institutional Animal Care and Use Committee of Washington State University and conducted in accordance with National Research Council guidelines and regulations for experiments in live animals.

### Surgical procedures

#### EEG &EMG and cannulae implantation

Adult male and female mice [Ctrl (n = 12; n = 6 females) and G_q_^+/-^ (n = 10; n = 5 females); 10 – 14-weeks-old)] were anesthetized with isoflurane and stereotaxically implanted with 2 chronic guide cannulae (C315GS-5/SPC; P1 Technologies, Roanoke, VA, USA) in basal forebrain (from Bregma AP: 0.0 mm, ML: ±1.62 mm, DV: -5.0 mm) [65,66]. Four electroencephalographic (EEG) screw electrodes (AMS120/3; Antrin Miniature Specialties, Fallbrook, CA, USA) were also implanted contralaterally over frontal (2) and occipital (2) cortices, and two electromyographic (EMG) wire electrodes were implanted in the nuchal muscles as previously described [2,3]. EEG electrodes and guide cannulae were fixed to the skull with dental acrylic. Patency of the cannulae was maintained with indwelling dummy cannulae (C315DCS-5/SPC; P1 Technologies). Mice were allowed to recover from surgery for at least 7 d prior to habituation to the recording environment.

#### Telemeter and cannulae implantation

We assessed changes in core temperature and gross motor activity to ensure that DREADD activation did not produce abnormal changes in physiology or behavior that might indirectly impact sleep-wake expression and regulation. A separate group of adult male and female G_q_^+/-^ mice (n = 5; n = 3 female; 9 – 14-weeks-old) were anesthetized with isoflurane and implanted with a telemetry device (G2 E-mitter; STARR Life Sciences Corp., Oakmont, PA, USA) in the intraperitoneal cavity as previously described [2,3]. A suture was used to secure the telemeter to the abdominal musculature, and wound clips were used to close the skin. Mice were then implanted with 2 chronic guide cannulae in the BF as described above and capped with indwelling dummy cannulae. Two anchor screws (Antrin Miniature Specialties) were placed contralaterally over frontal and visual cortices. Guide cannulae and anchor screws were fixed to the skull with dental acrylic. Body weight, hydration, and fecal output were monitored daily for 8 d after surgery at which point wound clips were removed.

### Sleep and polysomnographic analyses

#### CNO experiments

After recovery from cannulae, EEG, and EMG implantation, mice were individually housed in polycarbonate recording cages and connected to a lightweight, flexible recording cable [2,3,67]. Mice habituated to the recording cable for at least 3 d prior to data collection. Once habituated, baseline (BL) EEG and EMG data were collected for 24 h starting at light onset while mice were left undisturbed. The next day, either clozapine N-oxide dihydrochloride (CNO; 0.36 mM, i.e., 0.003 mg/kg for a 25 g mouse; HB6149: Hello Bio Inc., Princeton, NJ, USA) or vehicle (saline) was injected intracranially [68] into the BF via guide cannulae (250 nl per cannulae at ∼50 nl/min) in freely behaving mice using an internal cannula (C315IS-5/SPC; P1 Technologies) attached to a Hamilton syringe by silicone tubing (2415500; Dow Corning Corporation, Midland, MI, USA). CNO and vehicle injections occurred during Zeitgeber time (ZT) 0 using a counterbalance schedule separating injections by at least 48 h. Mice were left undisturbed after each injection, and EEG &EMG data were collected for 24 h. This approach allowed us to make within-subject vehicle vs. CNO comparisons as well as Ctrl vs. G_q_^+/-^ between-subject comparisons.

#### Sleep deprivation and J60 control experiments

We performed two additional experiments in the same Ctrl and G_q_^+/-^ mice that received vehicle and CNO. We first measured the homeostatic response to 6 hours of sleep deprivation (SD) to determine if the mice had a normal homeostatic response to SD. This would also provide comparison data to changes following DREADD activation in the G_q_^+/-^ mice. At least 48 h after the CNO/vehicle injections, Ctrl and G_q_^+/-^ mice underwent 6 h SD via gentle handling starting at light onset as previously described [2,3,7]. SD via gentle handling involves arousing mice (e.g., tactile, auditory stimuli) when their EEG/EMG and/or behavior (e.g., posture, quiescence) is predictive or indicative of sleep. Mice were then left to recover undisturbed for 18 h. Post-SD data were compared (and/or normalized where applicable) to time-matched values from the BL day.

Second, we determined if a different DREADD ligand (JHU37160 dihydrochloride [J60]; HB6261; Hello Bio Inc.) reproduced the sleep-wake effects of CNO in G_q_^+/-^ mice. This was prudent because CNO, but not J60, metabolizes to clozapine which may influence sleep via indirect effects [23,30]. Following the SD experiment, mice were allowed at least 48 h of additional recovery and then were injected intracranially with 0.35 mM J60 (i.e., 0.003 mg/kg for a 25 g mouse) during ZT0 (250 nl per cannulae at ∼50 nl/min), and EEG &EMG data were recorded for 24 h. These data were compared to results obtained from vehicle treatments used in the CNO experiments.

#### Polysomnography and EEG data analyses

EEG and EMG data were collected using a GRASS 7 polygraph system (Natus Medical Incorporated, Pleasanton, CA, USA) via a lightweight recording cable. The signals were amplified, digitized, and processed at 128 Hz using VitalRecorder acquisition software (v3.0.0.0; SleepSign for Animal, Kissei Comtec Co., LTD, Nagano, Japan). EEG and EMG data were high- and low-pass filtered at 0.3 and 100 Hz and 10 and 100 Hz, respectively [2,3]. Wakefulness, NREMS, and REMS were determined from EEG and EMG data by visual inspection of the EEG waveform, EMG activity, and fast Fourier transform (FFT) using SleepSign for Animal (v3.3.8.1803; Kissei Comtec Co., Ltd.). Vigilance states were scored in 4-s epochs by an investigator blinded to experimental conditions [2,3]. These data were then used to calculate time-in-state, bout duration, bout frequency, and latency to state. For vehicle, CNO, and J60 injection days, only post-injection data was included in calculations. Time-in-state was expressed as a percentage of total recording time (TRT) in 2-h bins. Minimum bout lengths were defined as ≥ 7 consecutive epochs (≥ 28 s) for wakefulness and NREMS and ≥ 4 consecutive epochs (≥ 16 s) for REMS [2,69]. Frequency and duration of vigilance state bouts were shown as differences from a control condition by subtracting time-matched vehicle data from CNO or J60 data or time-matched BL data from SD data [2,67,70]. These differences were then expressed in 6-h bins. Latency to NREMS and REMS was calculated in two ways. First, we determined elapsed time post-injection or post-SD before a bout of average length which was calculated from the 24-h vehicle data (or 24-h BL data for SD). We also calculated the latency to the first 4-s epoch of NREMS and REMS— an analysis which makes no assumption about minimal bout duration.

The EEG was fast Fourier transformed to produce power spectra between 0 and 30 Hz in 0.5 Hz bins [2]. All spectral data were normalized to undisturbed BL EEG spectra. Each spectral bin was expressed as a percentage of the total power in BL wakefulness, NREMS, and REMS averaged across the three vigilance states for the entire 24-h BL period [2,3]. We defined delta (δ) as 0.5 – 4 Hz, low δ as 0.5 – 1.5 Hz, theta (θ) as 5 – 9 Hz, alpha (α) as 10 – 15 Hz, and beta (β) as 15 – 30 Hz. As not all mice were equally awake or asleep in any given time bin, we used the following rules for calculating mean changes in EEG spectra as described previously [25]. A mouse had to spend ≥ 5 min in wakefulness or NREMS or ≥ 1 min in REMS (per time bin) for that EEG data to be included in the mean state analyses. In addition, for statistical comparisons, at least 5 mice per condition had to contribute to EEG spectra measurements per time bin. EEG epochs with visually detected artifacts were excluded from spectral analyses. Similar rules were applied to hourly changes in NREM δ power—defined as mean FFT power in δ (0.5 – 4 Hz) or low δ (0.5 – 1.5 Hz) [2,3,7]. FFT power within either the δ band or low δ band for each time bin post-injection or post-SD recovery were normalized to the average NREM δ or low δ band value, respectively, from the last 4 h of the undisturbed BL light period (h9 – 12) and expressed as a percentage shown in 2-h bins (adapted from [71]).

### Measurement of core temperature and activity

We measured changes in core temperature and gross activity after CNO and J60 because if present, they might indirectly alter sleep. After post-operative recovery from telemeter and cannulae implantation, mice were individually housed in standard mouse cages and given at least 5 days to habituate to the recording environment. As described above, mice were injected with either CNO or vehicle during ZT0 using a counterbalanced design separating the injections by 72 h. Three days later, all mice were injected with J60 during ZT0 as described above. After each injection, mice were left undisturbed, and core body temperature and cage activity were monitored continuously for 24 h. Core body temperature (°C) and gross cage activity data (counts) were captured from the abdominal telemetry device, transmitted to an energizer/receiver (ER-4000; STARR Life Sciences Corp.), and recorded with VitalView software (v5.1; STARR Life Sciences Corp.). Data were collected every minute for 24-h post-injection and averaged across 2-h bins to determine 24-h diurnal/nocturnal patterns in response to vehicle, CNO, and J60.

### Immunohistochemistry

We used immunohistochemistry to 1) confirm astroglial-specific expression of the DREADDs and 2) verify ligand activation of DREADD in astrocytes via astroglial cFos expression. Brains were obtained from the same mice used for sleep phenotyping at least 4 days after J60 injections. Ctrl (n = 3, females = 2) and G_q_^+/-^ (n = 3, females = 2) mice were injected with 0.36 mM CNO bilaterally in the BF (250 nl per cannulae at ∼50 nl/min) during ZT0, as described, and left undisturbed for approximately 90 minutes. Mice were then transcardially perfused with 1x phosphate buffered saline (PBS) followed by 10% buffered formalin. Brains were extracted and immediately post-fixed in 10% buffered formalin overnight at 4°C. To protect against freezing artifacts, brains were transferred to 4°C 30% sucrose in 1x PBS for 24 – 48 h and subsequently frozen and stored at -80°C until processing.

Frozen brains were sectioned coronally at 30 µm on a Thermo-Fisher CryoStar NX50 cryostat in series of 6 and stored free-floating in cryoprotectant. For immunohistochemical staining, free-floating sections were first washed 3 × 10 min in 1x PBS and then incubated for 30 min in a blocking solution of 2% normal goat serum (S-1000-20; Vector Laboratories, Newark, CA, USA) and 0.1% Triton x-100 (T8787-50ML; Sigma-Aldrich, St. Louis, MO, USA). Tissue was then incubated for 35 – 40 h at 4°C on a gentle rocker with primary antibodies diluted in blocking solution. Primary antibodies were used to amplify native mCitrine signal (1:1000, polyclonal guinea pig anti-GFP; 132-004; Synaptic Systems, Göttingen, Germany) and to identify astrocytes (1:1000, monoclonal rabbit anti-S100β; ab52642; Abcam, Waltham, MA, USA; 1:1000, polyclonal chicken anti-GFAP; ab4674; Abcam), cFos expression (1:1000, monoclonal mouse anti-cFos; 309-cFOS; PhosphoSolutions, Aurora, CO, USA), and neurons (1:1000, monoclonal mouse anti-NeuN; RBFOX/NeuN (1B7); Novus Biologicals, Centennial, CO, USA). Sections were then incubated for 1 h at room temperature with the following Alexa Fluor fluorophore-conjugated secondary antibodies diluted 1:1000 in blocking solution to generate fluorescence contrast of the primary antibodies for confocal detection: goat anti-guinea pig 488 (for GFP; A11073; Thermo Fisher Scientific, Waltham, MA, USA), donkey anti-rabbit 594 (for S100β; ab150132; Abcam), goat anti-chicken IgGy 594 (for GFAP; A11042; Thermo Fisher Scientific), goat anti-mouse (IgG1) 647 (for cFos; A21240; Thermo Fisher Scientific), and goat anti-mouse (IgG1) 594 (for NeuN; A21125; Thermo Fisher Scientific). After contrasting, sections were washed three times in 1x PBS, mounted onto generic 50 mm x 70 mm glass slides using 24 mm x 50 mm #1.5 thickness Gold Seal 3422 coverslips (50-189-9137; Fisher Scientific, Hampton, NH, USA) and DAPI Fluoromount-G (17984-24; Electron Microscopy Sciences, Hatfield, PA, USA).

An inverted Leica Microsystems DMi8 laser scanning microscope (Wetzlar, Germany) was used for image capture. Briefly, to identify sections containing basal forebrain structures, slides containing multiple mounted sections were first tile imaged at 5x with polarized light using the “Navigator” function in Leica Application Suite X software. Stereotaxic coordinates of our sections were then identified based on visual comparison against an atlas [72]. Using the Navigator function, BF structures were then digitally circumscribed to define a region of interest and optically sliced to capture volumes for quantification (25 – 30 1-µm thick sections at 512 × 512 pixels/frame using 20x HC PL APO 0.75 NA CS2 objective, 1.5x digital zoom). We primarily focused on the medial septum and the vertical limb of the diagonal band of Broca. To quantify nuclear cFos expression in BF astrocytes we required 1) that the fluorescence signals from DAPI and cFos completely colocalize in the x-y plane in a max-projection and when viewing the volume, be completely colocalized along the z-axis by optical cross-section and 2) colocalization of DAPI^+^/cFos^+^ nuclei with S100β tertiary labeling. A single mean was calculated across animals by averaging the total number of double-labeled cFos^+^ S100β^+^ cells across all regions of interest (ROIs) counted (n = 27 Ctrl ROIs; n = 28 G_q_^+/-^ ROIs).

### Statistical analysis

Plots were generated using SigmaPlot (v11.0, Systat Software, Inc., San Jose CA, USA) and R (v4.1.1). SPSS for Windows 25 (IBM Corporation, Armonk, NY, USA) was used for statistical analyses. Data are shown as means ± standard error of the mean (SE). Normality of the data was determined via Shapiro–Wilk or Kolmogorov–Smirnov tests. Normally distributed data were assessed with parametric tests: paired t test, unpaired t tests, one sample t test, general linear model for repeated measures (RM). In cases when data were not normally distributed or there were missing cells, we used non-parametric tests: Wilcoxon signed-rank tests, Mann-Whitney U tests, and Kruskal-Wallis tests. Comparisons of cell counts were made with Mann-Whitney U tests. RM using time (hours) as the repeated measure and either genotype (Ctrl vs. G_q_^+/-^) as the between-subjects factor or treatment (vehicle vs. CNO or J60; BL vs. SD) as the within-subjects factor was used for comparisons of several sleep and physiological measurements. These included time-in-state, bout frequency, bout duration, BL vs. SD NREM EEG δ power, core body temperature, and cage activity. For post-injection data, time-in-state (as % TRT), bout frequency, and bout duration RM comparisons were made over all time intervals for the full 24-h period. BL vs. SD time-in-state and NREM δ power was assessed over the initial 6-h recovery period in the light phase (h7 – 12). RM was also used for comparisons of EEG power spectra using frequency (Hz) as the repeated measure and treatment (vehicle vs. CNO or J60) as a within-subjects factors over the full 0 – 30 Hz range, or, as indicated, over the δ (0.5 – 4 Hz) and θ (5 – 9 Hz) frequency bands. RM comparisons were tested for sphericity, and a Greenhouse-Geisser correction was applied when needed. Post-hoc pairwise comparisons with Sidak corrections were performed when there were significant interaction effects or main effects of genotype or treatment. Comparisons of vehicle vs. CNO or J60 on NREM δ power and low NREM δ power were made using Kruskal-Wallis tests. Sidak corrections were applied to post-hoc pairwise comparisons. One sample t tests were used to determine if changes in bout frequency or bout duration differed from 0. Paired t tests or Wilcoxon signed-rank tests were used as appropriate for comparisons of sleep latency between treatments (vehicle vs. CNO or J60). Unpaired t tests or Mann-Whitney U tests were used as appropriate for comparisons of sleep latency between genotypes (Ctrl vs. G_q_^+/-^). When possible, sex was entered as a between-subjects factor separately for Ctrl and G_q_^+/-^mice for the measures described above. An alpha level less than 0.05 was used to indicate significance. To facilitate readability, statistical results are provided as supporting information in S1 - 11 Tables.

## Acknowledgements

We thank Dr. Christine Muheim, Dr. Kristan Singletary, and Sabrina Koh for their technical assistance.

## Notes

### Competing Interest Statement

The authors have declared no competing interest.

## References

1. Rosenwasser AM, Turek FW. Neurobiology of circadian rhythm regulation. Sleep Med Clin. 2022;17: 141–150.

2. Ingiosi AM, Frank MG. Noradrenergic signaling in astrocytes influences mammalian sleep homeostasis. Clocks & Sleep. 2022;4: 332–345.

3. Ingiosi AM, Hayworth CR, Harvey DO, Singletary KG, Rempe MJ, Wisor JP, et al. A role for astroglial calcium in mammalian sleep and sleep regulation. Curr Biol. 2020: Accepted.

4. Ingiosi AM, Frank MG. Goodnight, astrocyte: Waking up to astroglial mechanisms in sleep. FEBS J. 2022.

5. Martin-Fernandez M, Jamison S, Robin LM, Zhao Z, Martin ED, Aguilar J, et al. Synapse-specific astrocyte gating of amygdala-related behavior. Nat Neurosci. 2017;20: 1540–1548.

6. Ding F, O’Donnell J, Thrane AS, Zeppenfeld D, Kang H, Xie L, et al. A1-adrenergic receptors mediate coordinated ca2+ signaling of cortical astrocytes in awake, behaving mice. Cell Calcium. 2013;54: 387–394.

7. Halassa MM, Florian C, Fellin T, Munoz JR, Lee SY, Abel T, et al. Astrocytic modulation of sleep homeostasis and cognitive consequences of sleep loss. Neuron. 2009;61: 213–219.

8. Kim J-H, Kim J-H, Cho Y-E, Baek M-C, Jung J-Y, Lee M-G, et al. Chronic sleep deprivationinduced proteome changes in astrocytes of the rat hypothalamus. Journal of Proteome Research. 2014;13: 4047–4061.

9. Choi IS, Kim JH, Jeong JY, Lee MG, Suk K, Jang IS. Astrocyte-derived adenosine excites sleeppromoting neurons in the ventrolateral preoptic nucleus: Astrocyte-neuron interactions in the regulation of sleep. Glia. 2022;70: 1864–1885.

10. Vaidyanathan TV, Collard M, Yokoyama S, Reitman ME, Poskanzer KE. Cortical astrocytes independently regulate sleep depth and duration via separate gpcr pathways. eLife. 2021;10: e63329.

11. Durkee CA, Covelo A, Lines J, Kofuji P, Aguilar J, Araque A. G(i/o) protein-coupled receptors inhibit neurons but activate astrocytes and stimulate gliotransmission. Glia. 2019;67: 1076–1093.

12. Shen W, Chen S, Liu Y, Han P, Ma T, Zeng L-H. Chemogenetic manipulation of astrocytic activity: Is it possible to reveal the roles of astrocytes? Biochemical Pharmacology.2021;186: 114457.

13. Chai H, Diaz-Castro B, Shigetomi E, Monte E, Octeau JC, Yu X, et al. Neural circuit-specialized astrocytes: Transcriptomic, proteomic, morphological, and functional evidence. Neuron. 2017;95: 531-549.e539.

14. Spampinato SF, Copani A, Nicoletti F, Sortino MA, Caraci F. Metabotropic glutamate receptors in glial cells: A new potential target for neuroprotection? Frontiers in Molecular Neuroscience. 2018;11.

15. Kárpáti A, Yoshikawa T, Naganuma F, Matsuzawa T, Kitano H, Yamada Y, et al. Histamine h1 receptor on astrocytes and neurons controls distinct aspects of mouse behaviour. Scientific Reports. 2019;9: 16451.

16. Cai P, Huang S-N, Lin Z-H, Wang Z, Liu R-F, Xiao W-H, et al. Regulation of wakefulness by astrocytes in the lateral hypothalamus. Neuropharmacology. 2022;221: 109275.

17. Yang C, Thankachan S, McCarley RW, Brown RE. The menagerie of the basal forebrain: How many (neural) species are there, what do they look like, how do they behave and who talks to whom? Curr Opin Neurobiol. 2017;44: 159–166.

18. Brown RE, Basheer R, McKenna JT, Strecker RE, McCarley RW. Control of sleep and wakefulness. Physiol Rev. 2012;92: 1087–1187.

19. Adamsky A, Kol A, Kreisel T, Doron A, Ozeri-Engelhard N, Melcer T, et al. Astrocytic activation generates de novo neuronal potentiation and memory enhancement. Cell. 2018;174: 59–71 e14.

20. Traut J, Mengual JP, Meijer EJ, McKillop LE, Alfonsa H, Hoerder-Suabedissen A, et al. Effects of clozapine-n-oxide and compound 21 on sleep in laboratory mice. bioRxiv. 2022: 2022.2002.2001.478652.

21. Varin C, Luppi PH, Fort P. Melanin-concentrating hormone-expressing neurons adjust slow-wave sleep dynamics to catalyze paradoxical (rem) sleep. Sleep. 2018;41.

22. MacLaren DA, Browne RW, Shaw JK, Krishnan Radhakrishnan S, Khare P, España RA, et al. Clozapine n-oxide administration produces behavioral fffects in long-evans rats: Implications for designing dreadd experiments. eNeuro. 2016;3.

23. Manvich DF, Webster KA, Foster SL, Farrell MS, Ritchie JC, Porter JH, et al. The dreadd agonist clozapine n-oxide (cno) is reverse-metabolized to clozapine and produces clozapine-like interoceptive stimulus effects in rats and mice. Scientific Reports. 2018;8: 3840.

24. Jendryka M, Palchaudhuri M, Ursu D, van der Veen B, Liss B, Kätzel D, et al. Pharmacokinetic and pharmacodynamic actions of clozapine-n-oxide, clozapine, and compound 21 in dreadd-based chemogenetics in mice. Sci Rep. 2019;9: 4522.

25. Franken P, Chollet D, Tafti M. The homeostatic regulation of sleep need is under genetic control. J Neurosci. 2001;21: 2610–2621.

26. Vyazovskiy VV, Tobler I. Theta activity in the waking eeg is a marker of sleep propensity in the rat. Brain Res. 2005;1050: 64–71.

27. Zhang J, Zhu Y, Zhan G, Fenik P, Panossian L, Wang MM, et al. Extended wakefulness: Compromised metabolics in and degeneration of locus ceruleus neurons. The Journal of Neuroscience. 2014;34: 4418–4431.

28. Huber R, Deboer T, Tobler I. Effects of sleep deprivation on sleep and sleep eeg in three mouse strains: Empirical data and simulations. Brain Research. 2000;857: 8–19.

29. Harding EC, Franks NP, Wisden W. Sleep and thermoregulation. Current Opinion in Physiology. 2020;15: 7–13.

30. Bonaventura J, Eldridge MAG, Hu F, Gomez JL, Sanchez-Soto M, Abramyan AM, et al. Highpotency ligands for dreadd imaging and activation in rodents and monkeys. Nature Communications. 2019;10: 4627.

31. Dispersyn G, Sauvet F, Drogou C, Ciret S, Gallopin T, Chennaoui M. Validation of total sleep deprivation model in mice. Sleep Medicine. 2013;14: e107.

32. Bellesi M, Pfister-Genskow M, Maret S, Keles S, Tononi G, Cirelli C. Effects of sleep and wake on oligodendrocytes and their precursors. J Neurosci. 2013;33: 14288–14300.

33. Franken P, Malafosse A, Tafti M. Genetic determinants of sleep regulation in inbred mice. Sleep. 1999;22: 155–169.

34. Greene RW, Frank MG. Slow wave activity during sleep: Functional and therapeutic implications. The Neuroscientist. 2010;16: 618–633.

35. Frank MG, Morrissette R, Heller HC. Effects of sleep deprivation in neonatal rats. Am J Physiol. 1998;275: R148–157.

36. Dumoulin Bridi MC, Aton SJ, Seibt J, Renouard L, Coleman T, Frank MG. Rapid eye movement sleep promotes cortical plasticity in the developing brain. Sci Adv. 2015;1: e1500105.

37. Kim Y, Bolortuya Y, Chen L, Basheer R, McCarley RW, Strecker RE. Decoupling of sleepiness from sleep time and intensity during chronic sleep restriction: Evidence for a role of the adenosine system. Sleep. 2012;35: 861–869.

38. Guillaumin MCC, McKillop LE, Cui N, Fisher SP, Foster RG, de Vos M, et al. Cortical regionspecific sleep homeostasis in mice: Effects of time of day and waking experience. Sleep. 2018;41.

39. Milinski L, Fisher SP, Cui N, McKillop LE, Blanco-Duque C, Ang G, et al. Waking experience modulates sleep need in mice. BMC Biol. 2021;19: 65.

40. Shaw PJ, Franken P. Perchance to dream: Solving the mystery of sleep through genetic analysis. Journal of Neurobiology. 2003;54: 179–202.

41. Mang GM, Franken P. Genetic dissection of sleep homeostasis. Curr Top Behav Neurosci. 2015;25: 25–63.

42. Bjorness TE, Kelly CL, Gao T, Poffenberger V, Greene RW. Control and function of the homeostatic sleep response by adenosine a1 receptors. J Neurosci. 2009;29: 1267–1276.

43. Wimmer ME, Cui R, Blackwell JM, Abel T. Cyclic amp response element-binding protein is required in excitatory neurons in the forebrain to sustain wakefulness. Sleep. 2020;44.

44. Pedersen NP, Ferrari L, Venner A, Wang JL, Abbott SBG, Vujovic N, et al. Supramammillary glutamate neurons are a key node of the arousal system. Nature Communications. 2017;8: 1405.

45. Jhaveri KA, Trammell RA, Toth LA. Effect of environmental temperature on sleep, locomotor activity, core body temperature and immune responses of c57bl/6j mice. Brain Behav Immun. 2007;21: 975–987.

46. Kozak W, Zheng H, Conn CA, Soszynski D, van der Ploeg LH, Kluger MJ. Thermal and behavioral effects of lipopolysaccharide and influenza in interleukin-1 beta-deficient mice. Am J Physiol. 1995;269: R969–977.

47. Oka T, Oka K, Kobayashi T, Sugimoto Y, Ichikawa A, Ushikubi F, et al. Characteristics of thermoregulatory and febrile responses in mice deficient in prostaglandin ep1 and ep3 receptors. J Physiol. 2003;551: 945–954.

48. Xu M, Chung S, Zhang S, Zhong P, Ma C, Chang WC, et al. Basal forebrain circuit for sleep-wake control. Nat Neurosci. 2015;18: 1641–1647.

49. Kalinchuk AV, Porkka-Heiskanen T, McCarley RW, Basheer R. Cholinergic neurons of the basal forebrain mediate biochemical and electrophysiological mechanisms underlying sleep homeostasis. European Journal of Neuroscience. 2015;41: 182–195.

50. Kalinchuk AV, McCarley RW, Stenberg D, Porkka-Heiskanen T, Basheer R. The role of cholinergic basal forebrain neurons in adenosine-mediated homeostatic control of sleep: Lessons from 192 igg-saporin lesions. Neuroscience. 2008;157: 238–253.

51. Peng W, Wu Z, Song K, Zhang S, Li Y, Xu M. Regulation of sleep homeostasis mediator adenosine by basal forebrain glutamatergic neurons. Science. 2020;369: eabb0556.

52. Anaclet C, Pedersen NP, Ferrari LL, Venner A, Bass CE, Arrigoni E, et al. Basal forebrain control of wakefulness and cortical rhythms. Nat Commun. 2015;6: 8744.

53. Iwai Y, Ozawa K, Yahagi K, Mishima T, Akther S, Vo CT, et al. Transient astrocytic gq signaling underlies remote memory enhancement. Frontiers in Neural Circuits. 2021;15.

54. Gordon GRJ, Iremonger KJ, Kantevari S, Ellis-Davies GCR, MacVicar BA, Bains JS. Astrocytemediated distributed plasticity at hypothalamic glutamate synapses. Neuron. 2009;64: 391–403.

55. Kofuji P, Araque A. G-protein-coupled receptors in astrocyte–neuron communication. Neuroscience. 2021;456: 71–84.

56. Yang C, Franciosi S, Brown R. Adenosine inhibits the excitatory synaptic inputs to basal forebrain cholinergic, gabaergic, and parvalbumin neurons in mice. Frontiers in Neurology. 2013;4.

57. Hawryluk JM, Ferrari LL, Keating SA, Arrigoni E. Adenosine inhibits glutamatergic input to basal forebrain cholinergic neurons. J Neurophysiol. 2012;107: 2769–2781.

58. Arrigoni E, Chamberlin NL, Saper CB, McCarley RW. Adenosine inhibits basal forebrain cholinergic and noncholinergic neurons in vitro. Neuroscience. 2006;140: 403–413.

59. Yang C, Larin A, McKenna JT, Jacobson KA, Winston S, Strecker RE, et al. Activation of basal forebrain purinergic p2 receptors promotes wakefulness in mice. Scientific Reports. 2018;8: 10730.

60. Devaraju P, Sun MY, Myers TL, Lauderdale K, Fiacco TA. Astrocytic group i mglur-dependent potentiation of astrocytic glutamate and potassium uptake. J Neurophysiol. 2013;109: 2404–2414.

61. Wang F, Smith NA, Xu Q, Fujita T, Baba A, Matsuda T, et al. Astrocytes modulate neural network activity by ca2+-dependent uptake of extracellular k+. Sci Signal. 2012;5: ra26.

62. Loaiza A, Porras OH, Barros LF. Glutamate triggers rapid glucose transport stimulation in astrocytes as evidenced by real-time confocal microscopy. The Journal of Neuroscience. 2003;23: 7337–7342.

63. Scofield MD, Boger HA, Smith RJ, Li H, Haydon PG, Kalivas PW. Gq-dreadd selectively initiates glial glutamate release and inhibits cue-induced cocaine seeking. Biol Psychiatry. 2015;78: 441–451.

64. Benington JH. Sleep homeostasis and the function of sleep. Sleep. 2000;23: 1–8.

65. McKenna JT, Thankachan S, Uygun DS, Shukla C, McNally JM, Schiffino FL, et al. Basal forebrain parvalbumin neurons mediate arousals from sleep induced by hypercarbia or auditory stimuli. Curr Biol. 2020;30: 2379–2385 e2374.

66. Zant JC, Kim T, Prokai L, Szarka S, McNally J, McKenna JT, et al. Cholinergic neurons in the basal forebrain promote wakefulness by actions on neighboring non-cholinergic neurons: An opto-dialysis study. J Neurosci. 2016;36: 2057–2067.

67. Ingiosi AM, Schoch H, Wintler T, Singletary KG, Righelli D, Roser LG, et al. Shank3 modulates sleep and expression of circadian transcription factors. eLife. 2019;8: e42819.

68. Stachniak TJ, Ghosh A, Sternson SM. Chemogenetic synaptic silencing of neural circuits localizes a hypothalamus→midbrain pathway for feeding behavior. Neuron. 2014;82: 797–808.

69. Ingiosi AM, Hayworth CR, Harvey DO, Singletary KG, Rempe MJ, Wisor JP, et al. A role for astroglial calcium in mammalian sleep and sleep regulation. Current Biology. 2020;30: 4373-4383.e4377.

70. Ingiosi AM, Opp MR. Sleep and immunomodulatory responses to systemic lipopolysaccharide in mice selectively expressing interleukin-1 receptor 1 on neurons or astrocytes. Glia. 2016;64: 780–791.

71. Franken P, Tobler I, Borbely AA. Sleep homeostasis in the rat: Simulation of the time course of eeg slow-wave activity. Neurosci Lett. 1991;130: 141–144.

72. Franklin KBJ, Paxinos G (2008) The mouse brain in stereotaxic coordinates; Paxinos G, editor. Amsterdam: Elsevier.

