## Supporting information for "Activation of basal forebrain astrocytes induces wakefulness without compensatory changes in sleep drive"

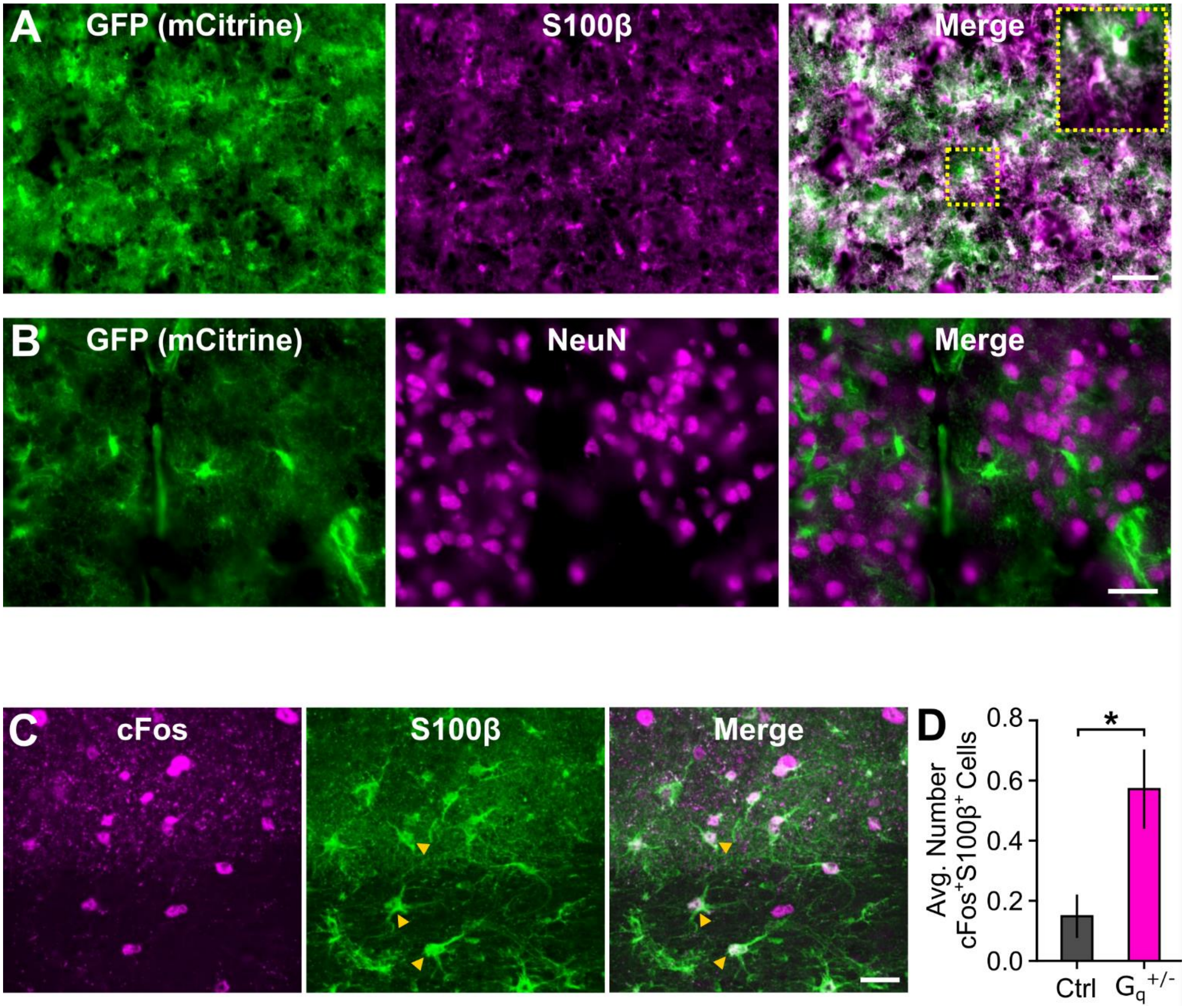


**S1 Fig: $G_q$-DREADDs are expressed in astrocytes of Aldh1l1-Cre$^{+/-}$ hM3Dq$^{fl/-}$ ($G_q^{+/-}$) mutant mice and CNO induces astroglial cFos expression in basal forebrain (BF) of $G_q^{+/-}$ mice.** (**A – B**) The surrogate $G_q$-DREADD mCitrine reporter amplified by an anti-GFP antibody colocalizes with the (**A**) astroglial marker S100β but not with the (**B**) neuronal maker NeuN in BF of $G_q^{+/-}$ mice. Scale bar for A is 50 µm (inset is 25 µm). Scale bar for B is 25 µm. (**C**) CNO-induced cFos expression in S100β-labeled astrocytes in $G_q^{+/-}$ BF. Examples of cFos$^+$S100β$^+$ double-labeled cells are marked with yellow arrowheads. Scale bar is 25 µm. (**D**) CNO increased cFos expression in BF astrocytes (labeled with S100β) in $G_q^{+/-}$ mice compared to CNO-treated control (Ctrl)

mice that do not express $G_q$-DREADDs. Values are means ± SE from n = 3 Ctrl mice (27 ROIs) and from n = 3 $G_q^{+/-}$ mice (28 ROIs; Mann-Whitney U, U = 252.5, * p = 0.009).

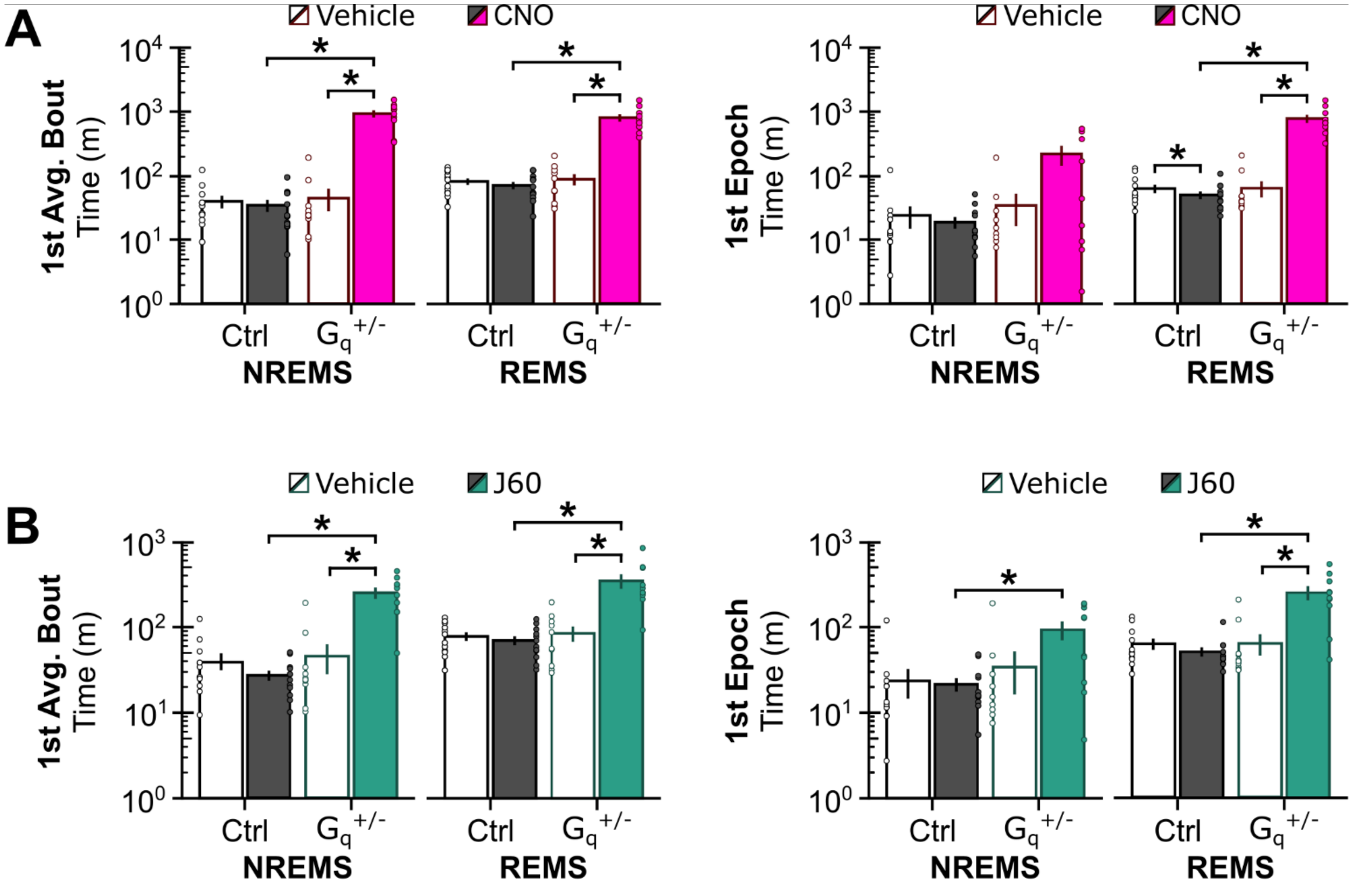


**S2 Fig: CNO and J60 increase latency to sleep in $G_q^{+/-}$ mice.** Latency to NREMS and REMS after (**A**) vehicle or CNO and (**B**) vehicle or J60 injections in BF defined as time to the first bout of average duration (based on 24-h vehicle data; left) or time to the first 4-s epoch (right) post-injection. Note the log10 scale on the y-axis (vehicle vs. CNO/J60: paired t test or Wilcoxon signed-rank; Ctrl vs. $G_q^{+/-}$: unpaired t test or Mann-Whitney U). Values are mean ± SE from n = 12 Ctrl and n = 10 $G_q^{+/-}$ mice. Dots are data from individual mice. * $p < 0.05$.

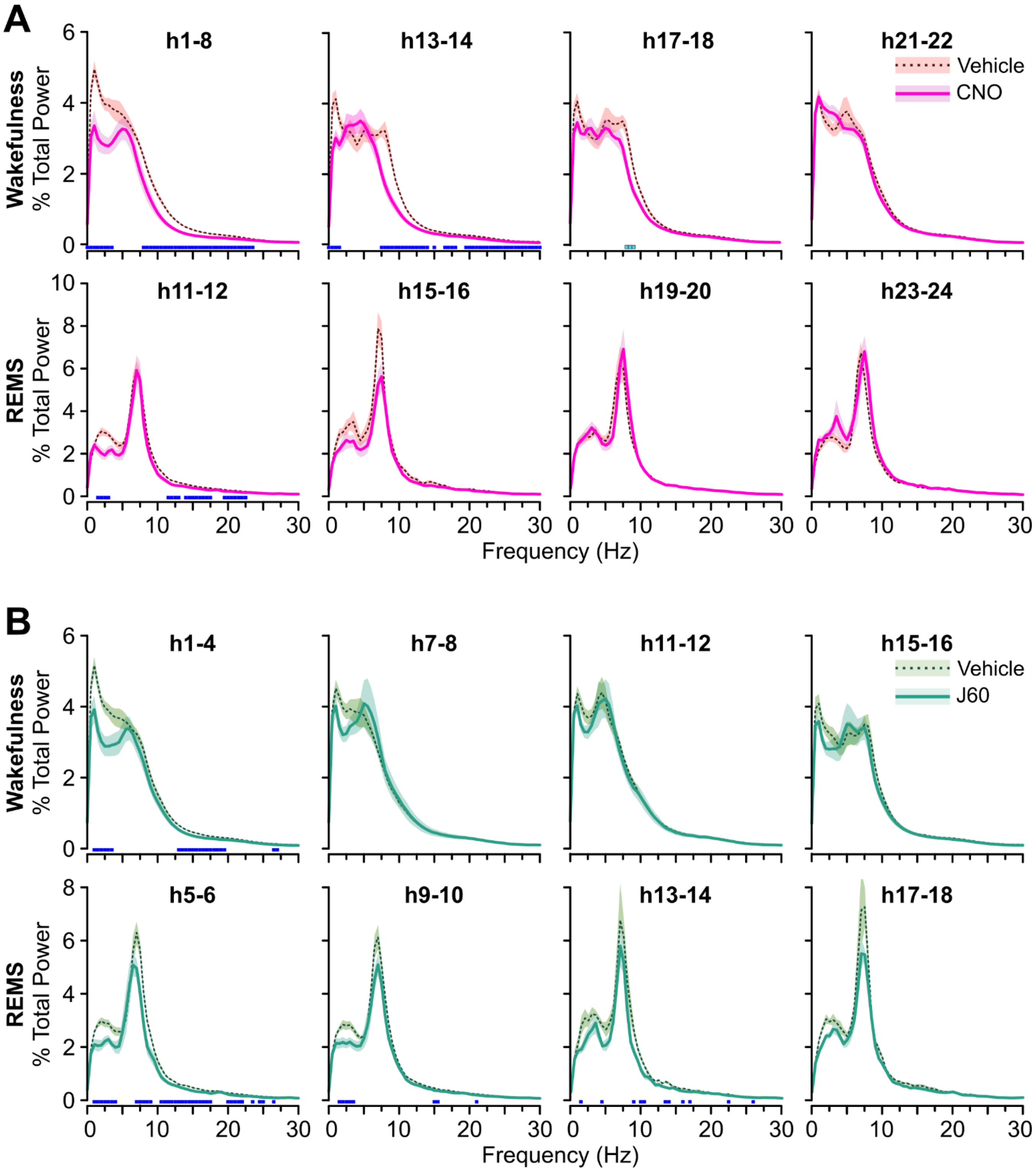


**S3 Fig: CNO and J60 reduce EEG spectral power in $G_q^{+/-}$ mice.** (**A**) Normalized EEG spectral power for wakefulness (top) and REMS (bottom) from $G_q^{+/-}$ mice shown in 2-h bins in response to vehicle and CNO. (**B**) Normalized EEG spectral power for wakefulness (top) and REMS (bottom) from $G_q^{+/-}$ mice shown in 2-h bins in response to vehicle and J60. Statistics are not shown for REMS h17-18 as only 4 mice had REMS after J60 during this time bin. For **A** and **B**, wakefulness plots start immediately after injection. REMS plots start from the first time bin at which ≥ 5 mice showed REMS post-injection. Light and dark blue squares above the x-axis denote frequency bins with significant vehicle vs. CNO/J60 differences (light blue: repeated measures ANOVA, 5 – 9 Hz; dark blue: repeated measures ANOVA, 0 – 30 Hz). Values are means (lines) ± SE (shading) based on data from n = 10 $G_q^{+/-}$ mice. $p < 0.05$.

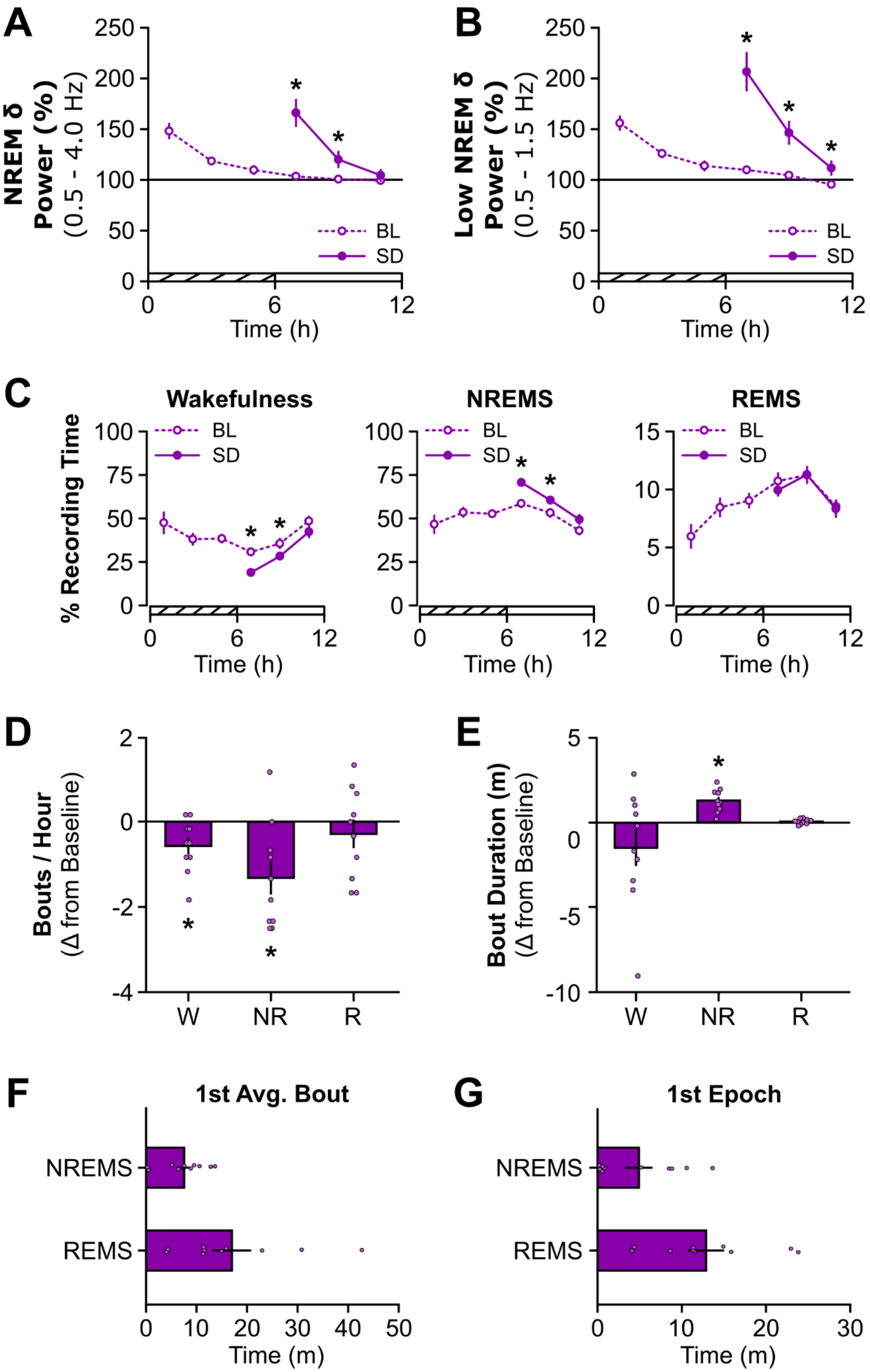
A
NREM δ Power (%) (0.5 - 4.0 Hz)
250
200
150
100
50
0
BL
SD
0
6
12
Time (h)
B
Low NREM δ Power (%) (0.5 - 1.5 Hz)
C
Wakefulness
NREMS
REMS
% Recording Time
100
75
25
15
10
5
D
Bouts / Hour (Δ from Baseline)
2
-2
-4
W
NR
R
E
Bout Duration (m) (Δ from Baseline)
-5
-10
F
1st Avg. Bout
NREMS
REMS
Time (m)
G
1st Epoch

**S4 Fig: Sleep deprivation increases sleep propensity in $G_q^{+/-}$ mice.** Normalized NREM delta (δ) power for the (**A**) full δ (0.5 – 4 Hz) and (**B**) low δ (0.5 – 1.5 Hz) bands shown in 2-h bins for undisturbed baseline (BL) sleep and recovery sleep after 6 h sleep deprivation (SD) for $G_q^{+/-}$ mice (repeated measures ANOVA). (**C**) Time spent in wakefulness (left), NREMS (middle), and REMS (right) under baseline conditions and after 6 h SD shown as a percentage of total recording time in 2-h bins for $G_q^{+/-}$ mice (repeated measures ANOVA). Cross-hatched bars on the x-axis for A - C denote the 6 h SD period. Open bars on the x-axis denote light period recovery phase. Change in (**D**) bout frequency and (**E**) bout duration shown as SD - BL differences for the first 6 h of recovery sleep (i.e., light period) post-SD. *, different from 0 (one-sample t test). Latency to the (**F**) first bout of average duration and (**G**) first epoch for NREMS and REMS after 6h SD. Values are means ± SE from n = 10 $G_q^{+/-}$ mice. Dots in D – G are data from individual mice. * $p < 0.05$.

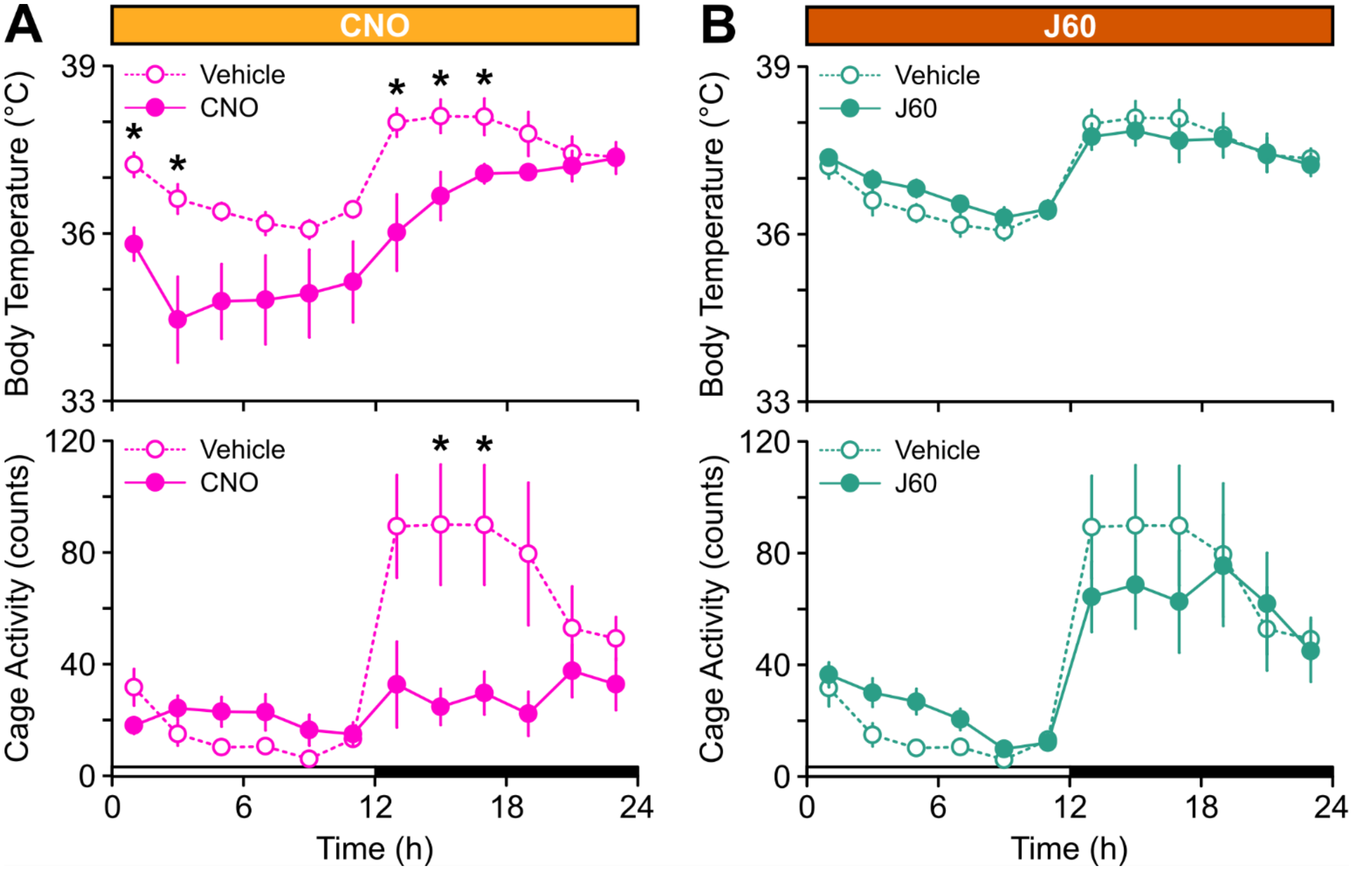


**S5 Fig: CNO, but not J60, reduces core body temperature and cage activity in $G_q^{+/-}$ mice.** (**A**) Core body temperature (top) and cage activity (bottom) for $G_q^{+/-}$ mice shown in 2-h bins in response to vehicle and CNO injected during ZT0. (**B**) Core body temperature (top) and cage activity (bottom) for $G_q^{+/-}$ mice shown in 2-h bins in response to vehicle and J60 injected during ZT0. Data are shown as means ± SE (repeated measures ANOVA). Open and closed bars on the x-axis represent the light and dark periods, respectively. n = 5 $G_q^{+/-}$ mice. * $p < 0.05$.

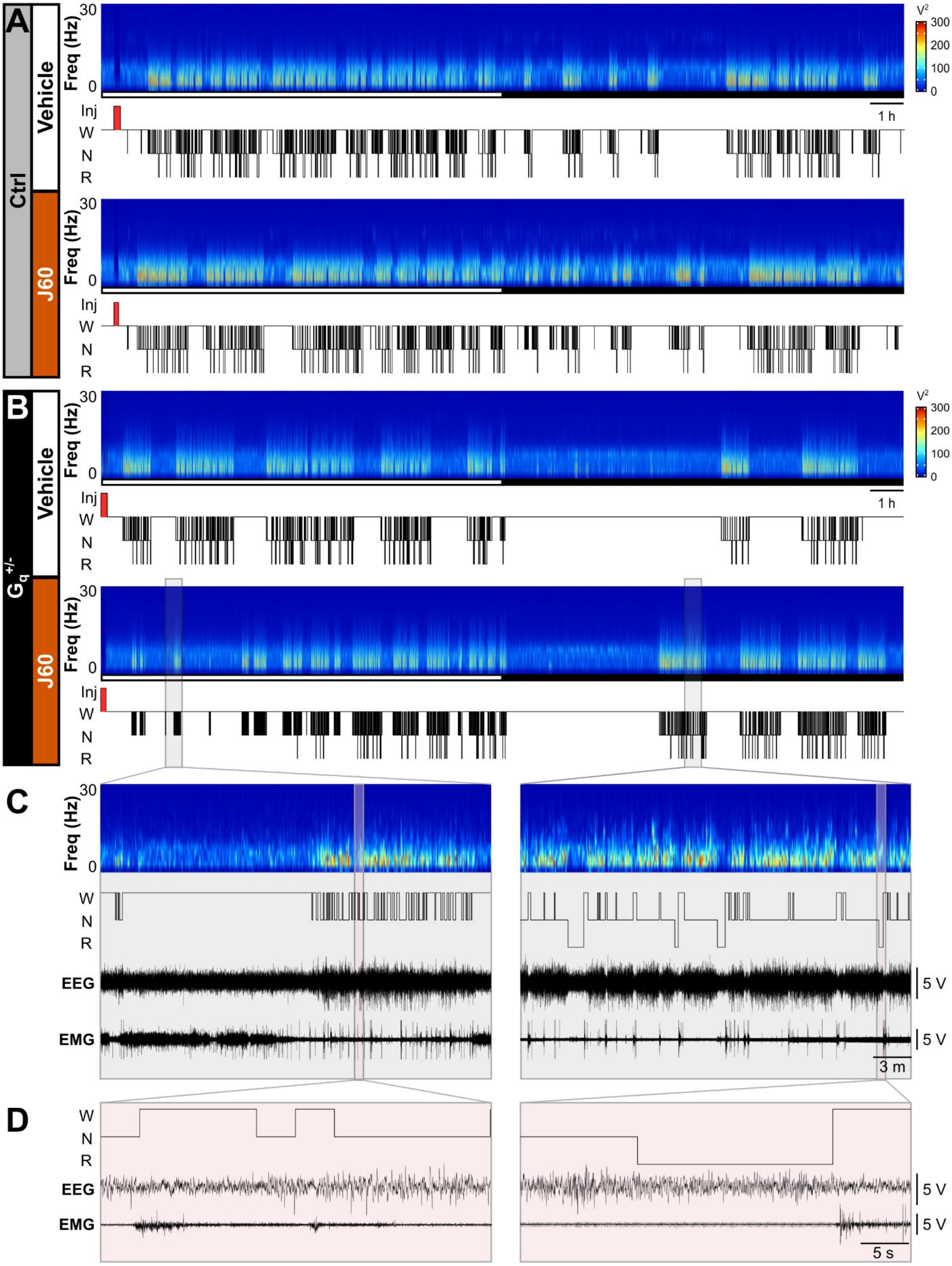
A
Ctrl
Vehicle
J60
Freq (Hz)
30
0
V²
300
200
100
0
1 h
Inj
W
N
R
B
$G_q^{+/-}$
C
D
EEG
EMG
5 V
3 m
5 s

**S6 Fig: J60 activation of $G_q$-DREADDs in basal forebrain astrocytes promotes wakefulness.**

Representative spectrograms, hypnograms, and EEG & EMG traces show responses to vehicle or J60 injections to the BF from the same (**A**) Ctrl and (**B**) $G_q^{+/-}$ mice shown in Fig 1. Open and closed bars below spectrograms represent the light and dark periods, respectively. (**C**) Gray boxes show 30-min subsets of $G_q^{+/-}$ post-J60 injection data from **B** for lower power, 'fragmented' sleep (left) and 'recovered' sleep (right). (**D**) Red boxes show 40-s subsets of $G_q^{+/-}$ post-J60 injection data extracted from the gray boxes in **C**. Injection timepoints are highlight by red rectangles in the hypnogram. Voltages are x$10^5$. Freq, frequency; Inj, injection; W, wakefulness; N, NREMS; R, REMS.

| | | CNO | | | | J60 | | | |
|---|---|---|---|---|---|---|---|---|---|
| | | numDF | denDF | F-value | p-value | numDF | denDF | F-value | p-value |
| **Time in state** | | | | | | | | | |
| | | *Ctrl: Vehicle vs. CNO* | | | | *Ctrl: Vehicle vs. J60* | | | |
| **Wakefulness** | Treatment | 1 | 22 | 0.14 | 0.716 | 1 | 22 | 4.08 | 0.056 |
| | Time x Treatment | 4.84 | 106.37 | 0.51 | 0.766 | 5.43 | 119.43 | 0.97 | 0.441 |
| **NREMS** | Treatment | 1 | 22 | 0.10 | 0.756 | 1 | 22 | 4.30 | 0.050 |
| | Time x Treatment | 4.83 | 106.22 | 0.49 | 0.776 | 5.42 | 119.30 | 0.95 | 0.459 |
| **REMS** | Treatment | 1 | 22 | 0.12 | 0.733 | 1 | 22 | 0.04 | 0.851 |
| | Time x Treatment | 5.03 | 110.73 | 0.51 | 0.768 | 4.49 | 98.81 | 0.83 | 0.520 |
| | | *$G_q^{+/-}$: Vehicle vs. CNO* | | | | *$G_q^{+/-}$: Vehicle vs. J60* | | | |
| **Wakefulness** | Treatment | 1 | 18 | 58.45 | **<0.001** | 1 | 18 | 8.78 | **0.008** |
| | Time x Treatment | 4.34 | 78.17 | 15.68 | **<0.001** | 4.28 | 76.96 | 5.40 | **<0.001** |
| **NREMS** | Treatment | 1 | 18 | 56.69 | **<0.001** | 1 | 18 | 7.81 | **0.012** |
| | Time x Treatment | 4.38 | 78.84 | 14.90 | **<0.001** | 4.17 | 75.07 | 5.21 | **<0.001** |
| **REMS** | Treatment | 1 | 18 | 32.54 | **<0.001** | 1 | 18 | 9.12 | **0.007** |
| | Time x Treatment | 4.39 | 78.96 | 17.23 | **<0.001** | 5.01 | 90.25 | 5.70 | **<0.001** |
| | | *Ctrl vs. $G_q^{+/-}$* | | | | *Ctrl vs. $G_q^{+/-}$* | | | |
| **Wakefulness** | Genotype | 1 | 20 | 82.24 | **<0.001** | 1 | 20 | 28.88 | **<0.001** |
| | Time x Genotype | 4.06 | 81.20 | 25.39 | **<0.001** | 5.46 | 109.20 | 9.38 | **<0.001** |
| **NREMS** | Genotype | 1 | 20 | 81.40 | **<0.001** | 1 | 20 | 27.44 | **<0.001** |
| | Time x Genotype | 4.08 | 81.50 | 22.95 | **<0.001** | 5.50 | 110.01 | 8.38 | **<0.001** |
| **REMS** | Genotype | 1 | 20 | 36.62 | **<0.001** | 1 | 20 | 16.05 | **<0.001** |
| | Time x Genotype | 4.30 | 86.04 | 30.09 | **<0.001** | 4.98 | 99.51 | 11.15 | **<0.001** |
| **Average Number Bouts / Hour** | | | | | | | | | |
| **Wakefulness** | Genotype | 1 | 20 | 8.94 | **0.007** | 1 | 20 | 5.07 | **0.036** |
| | Time x Genotype | 2.35 | 46.93 | 9.05 | **<0.001** | 3 | 60 | 0.48 | 0.695 |
| **NREMS** | Genotype | 1 | 20 | 10.92 | **0.004** | 1 | 20 | 0.25 | 0.626 |
| | Time x Genotype | 3 | 60 | 26.32 | **<0.001** | 3 | 60 | 3.79 | **0.015** |
| **REMS** | Genotype | 1 | 20 | 21.29 | **<0.001** | 1 | 20 | 4.99 | **0.037** |
| | Time x Genotype | 3 | 60 | 28.32 | **<0.001** | 3 | 60 | 11.97 | **<0.001** |
| **Average Bout Duration (min)** | | | | | | | | | |
| **Wakefulness** | Genotype | 1 | 20 | 4.65 | **0.043** | 1 | 20 | 0.27 | 0.612 |
| | Time x Genotype | 2.21 | 44.16 | 27.19 | **<0.001** | 2.10 | 41.97 | 4.97 | **0.011** |
| **NREMS** | Genotype | 1 | 20 | 106.64 | **<0.001** | 1 | 20 | 17.39 | **<0.001** |
| | Time x Genotype | 3 | 60 | 6.18 | **<0.001** | 2.17 | 43.49 | 3.95 | **0.024** |
| **REMS** | Genotype | 1 | 20 | 37.63 | **<0.001** | 1 | 20 | 19.46 | **<0.001** |
| | Time x Genotype | 3 | 60 | 11.64 | **<0.001** | 2.09 | 41.87 | 2.87 | 0.065 |

**S1 Table: Statistical output from repeated measures ANOVA comparisons of time in state, average number of bouts per hour, and average bout duration.**

| | | | Statistic | p-value |
|---|---|---|---|---|
| | | | **CNO** | |
| **Latency to NREM** | | | | |
| *Avg. bout duration* | Vehicle vs. CNO | Ctrl | Z = -1.02 | 0.308 |
| | | $G_q^{+/-}$ | Z = -2.80 | **0.005** |
| | Ctrl vs. $G_q^{+/-}$ | CNO | U = 0.00 | **<0.001** |
| *1st epoch* | Vehicle vs. CNO | Ctrl | Z = -0.16 | 0.875 |
| | | $G_q^{+/-}$ | Z = -1.68 | 0.093 |
| | Ctrl vs. $G_q^{+/-}$ | CNO | U = 38.00 | 0.159 |
| **Latency to REM** | | | | |
| *Avg. bout duration* | Vehicle vs. CNO | Ctrl | t(11) = 1.152 | 0.274 |
| | | $G_q^{+/-}$ | t(9) = -6.18 | **<0.001** |
| | Ctrl vs. $G_q^{+/-}$ | CNO | t(9.13) = -6.72 | **<0.001** |
| *1st epoch* | Vehicle vs. CNO | Ctrl | Z = -1.96 | **0.0497** |
| | | $G_q^{+/-}$ | Z = -2.80 | **0.005** |
| | Ctrl vs. $G_q^{+/-}$ | CNO | U = 0.00 | **<0.001** |
| | | | **J60** | |
| **Latency to NREM** | | | | |
| *Avg. bout duration* | Vehicle vs. J60 | Ctrl | Z = -1.33 | 0.182 |
| | | $G_q^{+/-}$ | Z = -2.80 | **0.005** |
| | Ctrl vs. $G_q^{+/-}$ | J60 | U = 1.00 | **<0.001** |
| *1st epoch* | Vehicle vs. J60 | Ctrl | Z = -1.02 | 0.308 |
| | | $G_q^{+/-}$ | Z = -1.78 | 0.074 |
| | Ctrl vs. $G_q^{+/-}$ | J60 | U = 25.00 | **0.021** |
| **Latency to REM** | | | | |
| *Avg. bout duration* | Vehicle vs. J60 | Ctrl | t(11) = 0.94 | 0.367 |
| | | $G_q^{+/-}$ | t(9) = -3.50 | **0.007** |
| | Ctrl vs. $G_q^{+/-}$ | J60 | t(9.30) = -4.07 | **0.003** |
| *1st epoch* | Vehicle vs. J60 | Ctrl | Z = -1.02 | 0.308 |
| | | $G_q^{+/-}$ | Z = -2.80 | **0.005** |
| | Ctrl vs. $G_q^{+/-}$ | J60 | U = 10.00 | **<0.001** |

**S2 Table: Statistical output for latencies to NREMS and REMS.**

| | CNO | | | J60 | | |
|---|---|---|---|---|---|---|
| | df | $\chi 2$ | p-value | df | $\chi 2$ | p-value |
| | | **Ctrl** | | | **Ctrl** | |
| **NREM Delta Power (0.5 - 4 Hz)** | 1 | 1.42 | 0.233 | 1 | 3.50 | 0.174 |
| **Low NREM Delta Power (0.5 - 1.5 Hz)** | 1 | 0.12 | 0.730 | 1 | 10.20 | **0.006** |
| | | **Gq+/-** | | | **Gq+/-** | |
| **NREM Delta Power (0.5 - 4 Hz)** | 1 | 7.78 | **0.005** | 1 | 10.61 | **0.005** |
| **Low NREM Delta Power (0.5 - 1.5 Hz)** | 1 | 30.42 | **<0.001** | 1 | 33.28 | **<0.001** |

**S3 Table: Statistical output from Kruskal Wallis comparisons of vehicle vs. CNO or J60 for NREM delta power and low NREM delta power.**

| | | | CNO | | | |
|---|---|---|---|---|---|---|
| | | | numDF | denDF | F-value | p-value |
| | | | Gq+/-: Vehicle vs. CNO | | | |
| **Wakefulness** | | | | | | |
| *h1 - 8* | 0 - 30 Hz | Treatment | 1 | 18 | 9.23 | **0.007** |
| | | Frequency x Treatment | 2.60 | 46.74 | 5.26 | **0.005** |
| *h13 - 14* | 0 - 30 Hz | Treatment | 1 | 18 | 5.51 | **0.031** |
| | | Frequency x Treatment | 3.52 | 63.44 | 5.17 | **0.002** |
| *h17 - 18* | 0 - 30 Hz | Treatment | 1 | 18 | 3.11 | 0.095 |
| | | Frequency x Treatment | 2.81 | 50.64 | 1.95 | 0.136 |
| | 5 - 9 Hz | Treatment | 1 | 18 | 5.05 | **0.037** |
| | | Frequency x Treatment | 1.38 | 24.84 | 1.20 | 0.301 |
| *h21 - 22* | 0 - 30 Hz | Treatment | 1 | 18 | 0.25 | 0.627 |
| | | Frequency x Treatment | 3.54 | 63.73 | 1.04 | 0.391 |
| **NREM** | | | | | | |
| *h9 - 10* | 0 - 30 Hz | Treatment | 1 | 14 | 3.95 | 0.067 |
| | | Frequency x Treatment | 2.30 | 32.22 | 3.17 | **0.049** |
| *h13 - 14* | 0 - 30 Hz | Treatment | 1 | 12 | 4.60 | 0.053 |
| | | Frequency x Treatment | 2.39 | 28.71 | 2.61 | 0.082 |
| | 0.5 - 4 Hz | Treatment | 1 | 12 | 3.09 | 0.104 |
| | | Frequency x Treatment | 2.68 | 32.17 | 4.43 | **0.013** |
| *h17 - 18* | 0 - 30 Hz | Treatment | 1 | 13 | 0.95 | 0.348 |
| | | Frequency x Treatment | 2.23 | 28.96 | 1.02 | 0.382 |
| *h21 - 22* | 0 - 30 Hz | Treatment | 1 | 17 | 1.44 | 0.247 |
| | | Frequency x Treatment | 2.00 | 33.93 | 1.24 | 0.303 |
| **REM** | | | | | | |
| *h11 - 12* | 0 - 30 Hz | Treatment | 1 | 13 | 4.89 | **0.046** |
| | | Frequency x Treatment | 3.20 | 41.62 | 1.19 | 0.326 |
| *h15 - 16* | 0 - 30 Hz | Treatment | 1 | 8 | 4.85 | 0.059 |
| | | Frequency x Treatment | 4.01 | 32.04 | 2.42 | 0.069 |
| *h19 - 20* | 0 - 30 Hz | Treatment | 1 | 14 | 0.01 | 0.947 |
| | | Frequency x Treatment | 2.92 | 40.82 | 1.22 | 0.313 |
| *h23 - 24* | 0 - 30 Hz | Treatment | 1 | 15 | 2.14 | 0.164 |
| | | Frequency x Treatment | 3.18 | 47.63 | 1.47 | 0.234 |

**S4 Table: Statistical output from repeated measures ANOVA comparisons of vehicle vs. CNO for $G_q^{+/-}$ EEG spectra.**

| | | Gq+/-: BL vs. SD | | | |
|---|---|---|---|---|---|
| | | numDF | denDF | F-value | p-value |
| | | **NREM delta power** | | | |
| **NREM Delta Power (0.5 - 4 Hz)** | | | | | |
| *Hours 7 - 12* | Treatment | 1 | 18 | 11.61 | **0.003** |
| | Time x Treatment | 1.11 | 19.92 | 43.81 | **<0.001** |
| **Low NREM Delta Power (0.5 - 1.5 Hz)** | | | | | |
| *Hours 7 - 12* | Treatment | 1 | 18 | 19.12 | **<0.001** |
| | Time x Genotype | 1.23 | 22.2 | 33.75 | **<0.001** |
| | | **Time in state** | | | |
| **Wakefulness** | | | | | |
| *Hours 7 - 12* | Treatment | 1 | 18 | 22.92 | **<0.001** |
| | Time x Treatment | 2 | 36 | 0.79 | 0.461 |
| **NREM** | | | | | |
| *Hours 7 - 12* | Treatment | 1 | 18 | 23.34 | **<0.001** |
| | Time x Treatment | 2 | 36 | 1.09 | 0.346 |
| **REM** | | | | | |
| *Hours 7 - 12* | Treatment | 1 | 18 | 0.3 | 0.591 |
| | Time x Treatment | 2 | 36 | 0.29 | 0.748 |

**S5 Table: Statistical output from repeated measures ANOVA comparisons of baseline (BL) vs. 6 h sleep deprivation (SD) for NREM delta power and time in state.**

| | | CNO | | | | J60 | | | |
|---|---|---|---|---|---|---|---|---|---|
| | | numDF | denDF | F-value | p-value | numDF | denDF | F-value | p-value |
| **Core Body Temperature** | Treatment | 1 | 8 | 7.60 | **0.025** | 1 | 8 | 0.11 | 0.750 |
| | Time x Treatment | 1.71 | 13.68 | 1.33 | 0.292 | 1.74 | 13.91 | 0.54 | 0.569 |
| **Cage Activity** | Treatment | 1 | 8 | 4.40 | 0.069 | 1 | 8 | 0.03 | 0.872 |
| | Time x Treatment | 2.09 | 16.71 | 4.08 | **0.035** | 1.58 | 12.62 | 0.79 | 0.448 |

**S6 Table: Statistical output from repeated measures ANOVA comparisons of vehicle vs. CNO or J60 for core body temperature and cage activity for $G_q^{+/-}$ mice.**

| | | | J60 | | | |
|---|---|---|---|---|---|---|
| | | | numDF | denDF | F-value | p-value |
| | | | **Gq+/-: Vehicle vs. J60** | | | |
| **Wakefulness** | | | | | | |
| *h1 - 4* | 0 - 30 Hz | Treatment | 1 | 18 | 4.06 | 0.059 |
| | | Frequency x Treatment | 1.70 | 30.62 | 4.24 | **0.029** |
| *h7 - 8* | 0 - 30 Hz | Treatment | 1 | 18 | 0.01 | 0.913 |
| | | Frequency x Treatment | 1.79 | 32.28 | 1.04 | 0.359 |
| *h11 - 12* | 0 - 30 Hz | Treatment | 1 | 18 | 0.21 | 0.650 |
| | | Frequency x Treatment | 2.43 | 43.75 | 0.47 | 0.668 |
| *h15 - 16* | 0 - 30 Hz | Treatment | 1 | 18 | 0.39 | 0.539 |
| | | Frequency x Treatment | 2.20 | 39.52 | 0.73 | 0.502 |
| **NREM** | | | | | | |
| *h3 - 4* | 0 - 30 Hz | Treatment | 1 | 14 | 2.15 | 0.165 |
| | | Frequency x Treatment | 1.29 | 17.98 | 2.42 | 0.132 |
| | 0.5 - 4 Hz | Treatment | 1 | 14 | 1.33 | 0.268 |
| | | Frequency x Treatment | 1.95 | 27.32 | 7.94 | **0.002** |
| *h7 - 8* | 0 - 30 Hz | Treatment | 1 | 18 | 1.72 | 0.207 |
| | | Frequency x Treatment | 1.92 | 34.51 | 0.70 | 0.496 |
| *h11 - 12* | 0 - 30 Hz | Treatment | 1 | 18 | 5.05 | **0.037** |
| | | Frequency x Treatment | 2.33 | 41.99 | 2.39 | 0.096 |
| *h15 - 16* | 0 - 30 Hz | Treatment | 1 | 10 | 1.39 | 0.265 |
| | | Frequency x Treatment | 2.15 | 21.47 | 0.46 | 0.653 |
| **REM** | | | | | | |
| *h5 - 6* | 0 - 30 Hz | Treatment | 1 | 15 | 6.69 | **0.021** |
| | | Frequency x Treatment | 2.75 | 41.23 | 2.44 | 0.083 |
| *h9 - 10* | 0 - 30 Hz | Treatment | 1 | 17 | 4.67 | **0.045** |
| | | Frequency x Treatment | 2.97 | 50.43 | 2.07 | 0.117 |
| *h13 - 14* | 0 - 30 Hz | Treatment | 1 | 8 | 6.18 | **0.038** |
| | | Frequency x Treatment | 2.41 | 19.29 | 0.98 | 0.406 |
| *h17 - 18* | 0 - 30 Hz | Treatment | - | - | - | - |
| | | Frequency x Treatment | - | - | - | - |

**S7 Table: Statistical output from repeated measures ANOVA comparisons of vehicle vs. J60 for $G_q^{+/-}$ EEG spectra.**

| | | Vehicle | | | | CNO | | | | J60 | | | |
|---|---|---|---|---|---|---|---|---|---|---|---|---|---|
| | | numDF | denDF | F-value | p-value | numDF | denDF | F-value | p-value | numDF | denDF | F-value | p-value |
| | | **Ctrl** | | | | **Ctrl** | | | | **Ctrl** | | | |
| **Wakefulness** | Sex | 1 | 10 | 1.07 | 0.326 | 1 | 10 | 0.61 | 0.454 | 1 | 10 | 2.40 | 0.153 |
| | Time x Sex | 3.47 | 34.69 | 0.65 | 0.607 | 3.95 | 39.45 | 1.25 | 0.307 | 4.74 | 47.35 | 1.18 | 0.334 |
| **NREMS** | Sex | 1 | 10 | 1.34 | 0.273 | 1 | 10 | 1.43 | 0.260 | 1 | 10 | 2.79 | 0.126 |
| | Time x Sex | 3.62 | 36.15 | 0.49 | 0.727 | 3.96 | 39.55 | 1.11 | 0.365 | 4.49 | 44.85 | 0.89 | 0.487 |
| **REMS** | Sex | 1 | 10 | 0.03 | 0.867 | 1 | 10 | 0.72 | 0.416 | 1 | 10 | 0.10 | 0.756 |
| | Time x Sex | 3.79 | 37.90 | 2.24 | 0.086 | 4.66 | 46.56 | 2.71 | **0.034** | 4.76 | 47.59 | 4.84 | **0.001** |
| | | **$G_q^{+/-}$** | | | | **$G_q^{+/-}$** | | | | **$G_q^{+/-}$** | | | |
| **Wakefulness** | Sex | 1 | 8 | 1.15 | 0.315 | 1 | 8 | 0.62 | 0.453 | 1 | 8 | 0.85 | 0.384 |
| | Time x Sex | 2.65 | 21.22 | 0.55 | 0.632 | 2.54 | 20.35 | 0.42 | 0.712 | 3.84 | 30.69 | 0.40 | 0.803 |
| **NREMS** | Sex | 1 | 8 | 1.86 | 0.210 | 1 | 8 | 0.67 | 0.437 | 1 | 8 | 0.96 | 0.356 |
| | Time x Sex | 2.66 | 21.30 | 0.56 | 0.625 | 2.46 | 19.65 | 0.53 | 0.636 | 3.86 | 30.89 | 0.45 | 0.767 |
| **REMS** | Sex | 1 | 8 | 0.93 | 0.363 | 1 | 8 | 0.26 | 0.622 | 1 | 8 | 0.21 | 0.657 |
| | Time x Sex | 3.14 | 25.09 | 0.53 | 0.675 | 2.95 | 23.59 | 0.18 | 0.908 | 3.09 | 24.72 | 0.65 | 0.598 |

**S8 Table: Statistical output from repeated measures ANOVA comparisons of males vs. females for time in state after vehicle, CNO, and J60 injections in Ctrl and $G_q^{+/-}$ mice.**

| | | CNO | | | | | | | | J60 | | | | | | | |
|---|---|---|---|---|---|---|---|---|---|---|---|---|---|---|---|---|---|
| | | Ctrl | | | | $G_q^{+/-}$ | | | | Ctrl | | | | $G_q^{+/-}$ | | | |
| | | numDF | denDF | F-value | p-value | numDF | denDF | F-value | p-value | numDF | denDF | F-value | p-value | numDF | denDF | F-value | p-value |
| | | **Average Number Bouts / Hour** | | | | **Average Number Bouts / Hour** | | | | **Average Number Bouts / Hour** | | | | **Average Number Bouts / Hour** | | | |
| **Wakefulness** | Sex | 1 | 10 | 0.12 | 0.741 | 1 | 8 | 0.71 | 0.425 | 1 | 10 | 3.52 | 0.090 | 1 | 8 | 0.65 | 0.444 |
| | Time x Sex | 3 | 30 | 0.50 | 0.683 | 3 | 24 | 0.91 | 0.449 | 3 | 30 | 1.04 | 0.391 | 3 | 24 | 1.45 | 0.254 |
| **NREMS** | Sex | 1 | 10 | 0.26 | 0.622 | 1 | 8 | 0.04 | 0.839 | 1 | 10 | 0.44 | 0.523 | 1 | 8 | 0.01 | 0.908 |
| | Time x Sex | 3 | 30 | 0.48 | 0.700 | 3 | 24 | 0.47 | 0.707 | 3 | 30 | 0.35 | 0.790 | 3 | 24 | 0.51 | 0.679 |
| **REMS** | Sex | 1 | 10 | 1.30 | 0.280 | 1 | 8 | 0.43 | 0.532 | 1 | 10 | 0.97 | 0.348 | 1 | 8 | 0.02 | 0.880 |
| | Time x Sex | 3 | 30 | 1.69 | 0.190 | 3 | 24 | 0.33 | 0.807 | 3 | 30 | 1.20 | 0.326 | 3 | 24 | 0.57 | 0.638 |
| | | **Average Bout Duration (min)** | | | | **Average Bout Duration (min)** | | | | **Average Bout Duration (min)** | | | | **Average Bout Duration (min)** | | | |
| **Wakefulness** | Sex | 1 | 10 | 0.80 | 0.393 | 1 | 8 | 1.79 | 0.218 | 1 | 10 | 4.34 | 0.064 | 1 | 8 | 0.27 | 0.619 |
| | Time x Sex | 1.97 | 19.72 | 1.66 | 0.216 | 3 | 24 | 0.44 | 0.725 | 3 | 30 | 1.73 | 0.181 | 3 | 24 | 0.23 | 0.878 |
| **NREMS** | Sex | 1 | 10 | 16.68 | **0.002** | 1 | 8 | 0.05 | 0.831 | 1 | 10 | 1.30 | 0.281 | 1 | 8 | 0.00 | 0.982 |
| | Time x Sex | 3 | 30 | 0.19 | 0.900 | 3 | 24 | 0.35 | 0.787 | 3 | 30 | 0.18 | 0.910 | 1.97 | 15.73 | 1.08 | 0.377 |
| **REMS** | Sex | 1 | 10 | 0.62 | 0.449 | 1 | 8 | 0.03 | 0.858 | 1 | 10 | 0.69 | 0.425 | 1 | 8 | 0.51 | 0.495 |
| | Time x Sex | 3 | 30 | 0.06 | 0.982 | 3 | 24 | 0.40 | 0.755 | 3 | 30 | 2.19 | 0.110 | 3 | 24 | 0.63 | 0.601 |

**S9 Table: Statistical output from repeated measures ANOVA comparisons of males vs. females for changes (from vehicle) in number of bouts per hour and bout duration for CNO and J60 treatment in Ctrl and $G_q^{+/-}$ mice.**

| | | | U statistic | p-value |
|---|---|---|---|---|
| **Latency to NREM** | | | | |
| *Avg. bout duration* | Vehicle | Ctrl | 9 | 0.150 |
| | | $G_q^{+/-}$ | 11 | 0.754 |
| | CNO | Ctrl | 10 | 0.200 |
| | | $G_q^{+/-}$ | 12 | 0.917 |
| | J60 | Ctrl | 11 | 0.262 |
| | | $G_q^{+/-}$ | 11 | 0.753 |
| *1st epoch* | Vehicle | Ctrl | 11 | 0.262 |
| | | $G_q^{+/-}$ | 5 | 0.117 |
| | CNO | Ctrl | 13 | 0.423 |
| | | $G_q^{+/-}$ | 10 | 0.602 |
| | J60 | Ctrl | 14 | 0.522 |
| | | $G_q^{+/-}$ | 10 | 0.602 |
| **Latency to REM** | | | | |
| *Avg. bout duration* | Vehicle | Ctrl | 18 | 1.000 |
| | | $G_q^{+/-}$ | 7 | 0.251 |
| | CNO | Ctrl | 17 | 0.873 |
| | | $G_q^{+/-}$ | 11 | 0.754 |
| | J60 | Ctrl | 11 | 0.262 |
| | | $G_q^{+/-}$ | 11 | 0.251 |
| *1st epoch* | Vehicle | Ctrl | 10 | 0.200 |
| | | $G_q^{+/-}$ | 6 | 0.175 |
| | CNO | Ctrl | 14 | 0.522 |
| | | $G_q^{+/-}$ | 12 | 0.917 |
| | J60 | Ctrl | 11 | 0.262 |
| | | $G_q^{+/-}$ | 11 | 0.754 |

**S10 Table: Statistical output from Mann Whitney U comparisons of males vs. females for latency to NREMS and REMS after vehicle, CNO, and J60 injections in Ctrl and $G_q^{+/-}$ mice**

| | Vehicle | | | CNO | | | J60 | | |
|---|---|---|---|---|---|---|---|---|---|
| | df | $\chi^2$ | p-value | df | $\chi^2$ | p-value | df | $\chi^2$ | p-value |
| | | **Ctrl** | | | **Ctrl** | | | **Ctrl** | |
| **NREM Delta Power (0.5 - 4 Hz)** | 1 | 1.22 | 0.269 | 1 | 1.86 | 0.173 | 1 | 0.48 | 0.487 |
| **Low NREM Delta Power (0.5 - 1.5 Hz)** | 1 | 1.36 | 0.243 | 1 | 5.84 | **0.016** | 1 | 1.00 | 0.319 |
| | | **$G_q^{+/-}$** | | | **$G_q^{+/-}$** | | | **$G_q^{+/-}$** | |
| **NREM Delta Power (0.5 - 4 Hz)** | 1 | 7.38 | **0.007** | 1 | 7.16 | **0.007** | 1 | 3.82 | 0.051 |
| **Low NREM Delta Power (0.5 - 1.5 Hz)** | 1 | 0.05 | 0.832 | 1 | 3.56 | 0.059 | 1 | 3.73 | 0.053 |

**S11 Table: Statistical output from Kruskal Wallis comparisons of males vs. females for NREM delta power and low NREM delta power after vehicle, CNO, and J60 injections for Ctrl and $G_q^{+/-}$ mice.**
